# Cathepsin-dependent amyloid formation drives mechanical rupture of lysosomal membranes

**DOI:** 10.64898/2026.01.17.700056

**Authors:** Delong Li, Wenxin Zhang, Michaela Medina, Jan F. M. Stuke, Andre Schwarz, Jonas Brill, Johann Brenner, Felix Kraus, Simon Ohlerich, Javier Lizarrondo, Jeremy Pflaum, Julia H. Grass, Lena-Marie Soltow, Dietmar Hammerschmid, Natalie Weber, Sonja Welsch, Julian D. Langer, Maike Windbergs, J. Wade Harper, Erin Schuman, Gerhard Hummer, Danielle A. Grotjahn, Florian Wilfling

## Abstract

Lysosomal membrane integrity is essential for cellular homeostasis, and its failure drives lysosomal storage disorders (LSD) and neurodegeneration. The dipeptide L-leucyl-L-leucine methyl ester (LLOMe) is widely used to model lysosomal damage, yet its mechanism remains poorly understood. The prevailing view holds that LLOMe polymerizes into membrane-permeabilizing peptide chains within the lysosomal lumen. Using cryo-electron tomography in cultured cells and primary neurons, we visualized the structural basis of LLOMe-induced lysosomal damage. We reveal that LLOMe forms amyloid structures within lysosomes that directly interact with and rupture the limiting membrane through mechanical stress. *In vitro* reconstitution confirms this amyloid-mediated mechanism. These findings establish a structural paradigm for lysosomal membrane disruption and provide insights into how disease-relevant protein aggregates, implicated in neurodegeneration and LSD, may compromise lysosomal integrity.

## Introduction

Lysosomes serve as the cell’s primary degradative compartment, breaking down macromolecules and damaged organelles to maintain cellular homeostasis (*1*). The lysosomal membrane acts as a critical barrier, sequestering hydrolytic enzymes with optimal activity at acidic pH. Loss of this membrane integrity, termed lysosomal membrane permeabilization (LMP), can result in the release of hydrolytic contents into the cytosol, triggering cell death pathways (*2, 3*). In neurodegenerative diseases, lysosomal damage enables the escape of amyloid aggregates and fibrils into the cytoplasm, where they seed further pathological aggregation and disease progression (*4–7*). To counteract such damage, cells employ quality control mechanisms including membrane repair machinery and autophagy-mediated clearance of damaged lysosomes (*8–14*).

LMP is thought to manifest through various structural alterations, ranging from localized lipid packing defects and progressive membrane disintegrity to the formation of discrete pores or overt rupture (*15, 16*). Despite its pathological significance, the ultrastructural basis of lysosomal membrane failure remains poorly characterized.

The dipeptide L-leucyl-L-leucine methyl ester (LLOMe) has been used for four decades as a potent tool to induce lysosomal membrane permeabilization (*17, 18*). Upon entering lysosomes, the methyl ester of LLOMe is cleaved, and the resulting product is thought to polymerize into membrane-damaging species (*19, 20*). However, the mechanism by which this dipeptide methyl ester polymerizes and disrupts biological membranes remains unclear, and direct ultrastructural visualization of LLOMe-induced lysosomal damage has not been achieved.

## Results

### *In situ* lysosomal ultrastructure and identity

To investigate changes in lysosomal membrane ultrastructure upon LLOMe treatment, we employed 2D-correlative cryo-focused ion beam milling (cryo-FIB) and cryo-ET to first visualize lysosomal structures in HeLa cells under basal conditions as our reference state (Fig. S1A). Lysosomes were identified *in situ* using 10kDa Dextran coupled to Alexa Fluor 647 (Dextran-AF647), which after endocytic uptake accumulates in endo-lysosomal compartments following overnight incubation (*21, 22*). Cryo-ET tomogram reconstruction showed the previously reported electron-dense interior of lysosomal structures with multiple membrane whorls within these compartments (*23*), suggestive of ongoing phospholipid degradation (Fig. 1A). These lysosomal structures were as expected tightly integrated in the cellular organelle network including the ER, mitochondria, and nucleus. To quantify these contacts more precisely we measured the frequency of lysosome-organelle contacts (Fig. S1B) and their distances, which were found to be in average between 15–25 nm (Fig. S1C).

**Fig. 1.**
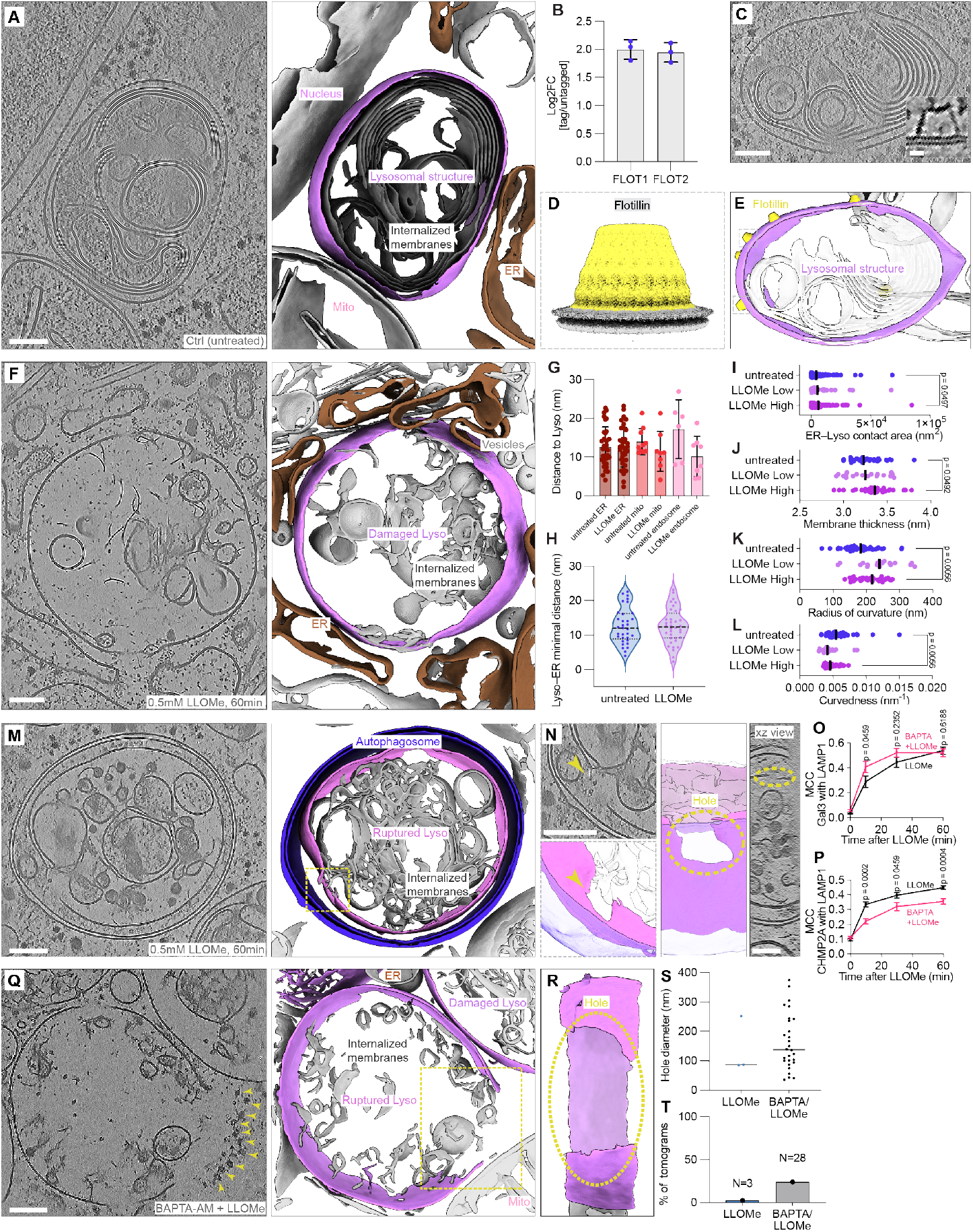
In situ cryo-ET reveals lysosomal ultrastructure and identity. (**A**) Tomographic slice (left) and corresponding segmentation (right) showing a lysosomal structure with its surrounding components (lysosome in purple, ER in brown). (**B**) Mean log2FC of Flotillin complex (FLOT1/FLOT2) in tagged/untagged LysoIP-MS plotted from (25). (**C**) Tomographic slice of a lysosomal structure with Flotillin complexes. Zoom-in area: a rotated view of the flotillin complex. (**D**) Sharpened Flotillin sub-tomogram averaging map, reaching 18.74Å. (**E**) Corresponding segmentation of (**C**) with Flotillin coordinates placed back into the tomogram. (**F**) Tomographic slice of a damaged lysosome after LLOMe treatment (0.5 mM, 1 h) (left) and corresponding segmentation (right). Lysosomal membrane in purple, surrounding ER in brown. (**G**) Minimal distance of lysosomal structure to various organelles (mean ± s.e.m.). (**H**) Minimal distance of lysosomal structure to ER. (**I**)-(**L**) ER-lysosome contact area, membrane thickness, local radius of curvature and global curvature calculated from lysosomal structures in untreated cells or in LLOMe-treated cells (0.5mM, 1 h) with low and high degrees of damage. (**M**) Tomographic slice (left) and corresponding segmentation (right) exhibiting an autophagosome encapsulating a ruptured lysosome upon LLOMe treatment (0.5mM, 1 h). The autophagosome membrane in blue, lysosomal membrane in purple. (**N**) Zoom-in area in tomogram and segmentation showing a hole (yellow arrow) in the delimiting lysosomal membrane. Side view of hole in lysosomal membrane, hole outlined using dashes. (**O**-**P**) Mander’s colocalization coefficient (MCC) quantifying the fraction of GAL-3 or CHMP2A overlapping with LAMP1 (n=12-16 fields, mean ± s.e.m.). Cells were treated with or without 50 μM BAPTA for 30min, followed by 0.5 mM LLOMe treatment for the indicated time points. (**Q**) Tomographic slice (left) and corresponding segmentation (right) showing a lysosome with clear membrane rupture after pre-treatment with 50 µM BAPTA (30 min), followed by 0.5 mM LLOMe for 1 h. Lysosomes in purple, ribosomes indicated by yellow arrowheads, a hole in the delimiting membrane of the central lysosome outlined by dashes. (**R**) Side view of the hole in panel Q. (**S**) Quantification of hole size in ruptured lysosomes (LLOMe, n=3, BAPTA +LLOMe n=28; median). (**T**) Quantification of the frequency of lysosomes containing holes across the entire datasets (LLOMe, n=115; BAPTA +LLOMe, n=116). Scale bars: 100 nm (**A, C, F, M, N** and **Q**), 10 nm (zoomed-in inset in **C**). Mann-Whitney test was conducted with p-values indicated (**I**-**L**, and **O**-**P**).

Flotillin-1 and Flotillin-2 have recently been identified as marker proteins that distinguish early endosomes from endo-lysosomal structures (*24*). To further confirm the identity of the structures classified as lysosomes based on morphology, we reanalyzed existing Lyso-IP proteomics data from our HeLa cells containing an endogenously tagged TMEM192-3xHA (*25*) and found that the flotillin complex, comprising Flotillin-1 and Flotillin-2, was significantly enriched in the tagged over the untagged sample (Fig. 1B, Fig. S1D). As the flotillin complex exclusively marks lysosomes (*24*) and exhibits a distinctive structure characteristic of SPFH family proteins (*26*), it allowed for visual identification in our tomograms (Fig. 1C). We performed templatematching and sub-tomogram averaging to determine the structure of the Flotillin complex *in situ* at 18.74 Å resolution (Fig. 1D, Fig. S1E-F), which allowed fitting of the published atomic model (Fig. S1G). Mapping the coordinates of the averaged Flotillin sub-tomograms back onto membrane segmentations confirmed that the Flotillin complex localizes to the surface of morphologically identified lysosomal structures (Fig. 1E).

### LLOMe triggers lysosomal membrane rupture

Having established the native lysosomal ultrastructure in HeLa cells, we next investigated changes in lysosomal membrane structure upon treatment with the lysosomotropic agent LLOMe. LLOMe treatment has been shown to induce several repair pathways, e.g., as recruitment of the ESCRT machinery (*12, 14, 27*). After loss of lysosomal membrane integrity, Galectin-3 (GAL-3), a β-galactoside-binding protein that binds damage-exposed glycans, colocalizes with lysosomal markers (*28, 29*).

We first examined the recruitment of GAL-3 and the ESCRT-III component CHMP2A to lysosomes upon LLOMe treatment (Fig. S2A-B). Consistent with previous reports (*27–29*), we observed an increased number of GAL-3 puncta and the progressive recruitment of GAL-3 and CHMP2A to LLOMe-treated lysosomes over time (Fig. S2C), confirming the kinetics of the damage response. Based on these kinetics, we predicted maximal lysosomal ultrastructural damage at 60 minutes of LLOMe treatment and acquired 115 cryo-ET tilt series of LLOMe-treated lysosomes in HeLa cells (Fig. 1F). We observed again the flotillin complex on lysosomal surfaces (Fig. S2D) and furthermore noticed structural evidence of three known lysosomal damage repair pathways: direct lipid transfer from the endoplasmic reticulum (ER) to lysosomes (*30, 31*), putative ATG9A vesicles of approximately 40 nm diameter (*32*), and membrane sealing likely mediated by ESCRT (*27*) (Fig. S2E). Notably, the ER extensively enveloped damaged lysosomes upon LLOMe-treatment (Fig. 1F), consistent with previous reports demonstrating that ER-mediated lipid synthesis and transfer are required for efficient membrane repair (*31, 33*). To quantify these changes, we performed morphometric analyses of lysosome-organelle interactions. Distances between lysosomes and other organelles, remained unchanged between untreated and LLOMe-treated cells (Fig. 1G), with minimal ER-lysosome distances of approximately 12 nm maintained regardless of treatment condition (Fig. 1H). These data suggest that contact-site machinery preserves these inter-organelle distances independent of lysosomal damage.

However, while contact distances remained constant, visual inspection revealed heterogeneous levels of membrane damage in LLOMe-treated lysosomes, prompting us to examine whether ER coverage scales with damage severity. We stratified the dataset into low and high degrees of lysosomal damage based on lysosome morphology and observed a significant increase in ER-lysosome contact area with higher damage severity (Fig. 1I). These findings indicate that ER wrapping of damaged lysosomes occurs in a controlled yet dynamic manner: contact-site distances are maintained by dedicated machinery, but the extent of ER coverage increases in proportion to damage severity.

Furthermore, we examined morphometrical changes of the lysosome upon LLOMe treatment: lysosomal membrane thickness increased modestly but significantly from 3.23 nm in untreated lysosomes to 3.36 nm in the highly damaged subset of LLOMe-treated lysosomes (Fig 1J), suggesting altered membrane composition in highly damaged lysosomes. Additionally, treated lysosomes exhibited a significant increase in the radius of curvature and inversely in curvature (Fig. 1K-L), indicating lysosomal swelling upon LLOMe treatment.

While most LLOMe-treated lysosomes maintained membrane continuity, we observed large holes in a subset of tomograms (Fig. S2F). In one instance, a lysosome with a hole of ∼150 nm diameter was encapsulated within an autophagosome (Fig. 1M-N). Such rupture events were rare, occurring in only approximately 3% of tomograms, likely reflecting both experimental constraints and efficient repair and degradation mechanisms (*14, 27, 31*).

To confirm that lysosomal membrane rapture was indeed induced by LLOMe treatment, we inhibited ESCRT-mediated membrane repair prior to LLOMe treatment by pretreating cells with the membrane-permeable calcium chelator BAPTA-AM. Ca^2+^ release is thought to be a key rapid response mechanism following lysosomal membrane disruption (*34*). Consistent with previous studies, BAPTAAM pretreatment accelerated GAL-3 recruitment to LLOMe treated lysosomes (Fig. 1O), with a concomitant reduction of CHMP2A recruitment (*27, 29*) (Fig. 1P, Fig. S3A-B). We acquired 116 cryo-ET tilt series from BAPTA-AMpretreated, LLOMe-treated cells and found ruptured lysosomes in 24% of tomograms (Fig. 1T, Fig. S3C). Hole diameters ranged from 50 nm to 400 nm (Fig. 1S). One extreme case displayed a ruptured lysosome with a hole diameter of approximately 450 nm (Fig. 1Q-R). Notably, this snapshot captured lysosomal contents extruding into the cytosol, displacing adjacent ribosomes from the immediate vicinity of the ruptured membrane (Fig. 1Q). Additional instances of autophagosome engulfment of damaged lysosomes were also observed (Fig. S3D-E). Collectively, these data demonstrate that LLOMe ruptures lysosomes by creating membrane holes.

### LLOMe forms fibrillar structures both in HeLa cells and rat primary neurons

LLOMe within the lysosomal lumen is expected to form small polymers induced by cathepsin C (CatC) (*19*). Surprisingly, we observed unexpected electron-dense structures within the lysosomal lumen following LLOMe treatment (Fig. 2A). Segmentation of these structures revealed that they form fibrillar to sheet-like assemblies (Fig. 2B-C). To determine whether these fibrillar structures are cell-type specific, we examined primary rat hippocampal neurons. Similar to HeLa cells, lysosomes were visualized using Dextran-AF647 labelling, and neurons were identified within the rat brain preparation using the live-neuron dye NeuO (*35*) (Fig. 2D). In untreated neurons, lysosomes exhibited a similar multi-membrane whorl architecture as observed in HeLa cells (Fig. 2E). Following 1 h of LLOMe treatment, neuronal lysosomes exhibited substantially larger diameters, forming vacuole-like structures visible by confocal live-cell microscopy (Fig. 2F). These enlarged lysosomal vacuoles contained massive accumulation of associated electron-dense structures which were evident in cryo-ET tomograms (Fig. 2G). Importantly, the structures exhibited similar fibrillar architectures as seen in HeLa cells (Fig. 2H)

**Fig. 2.**
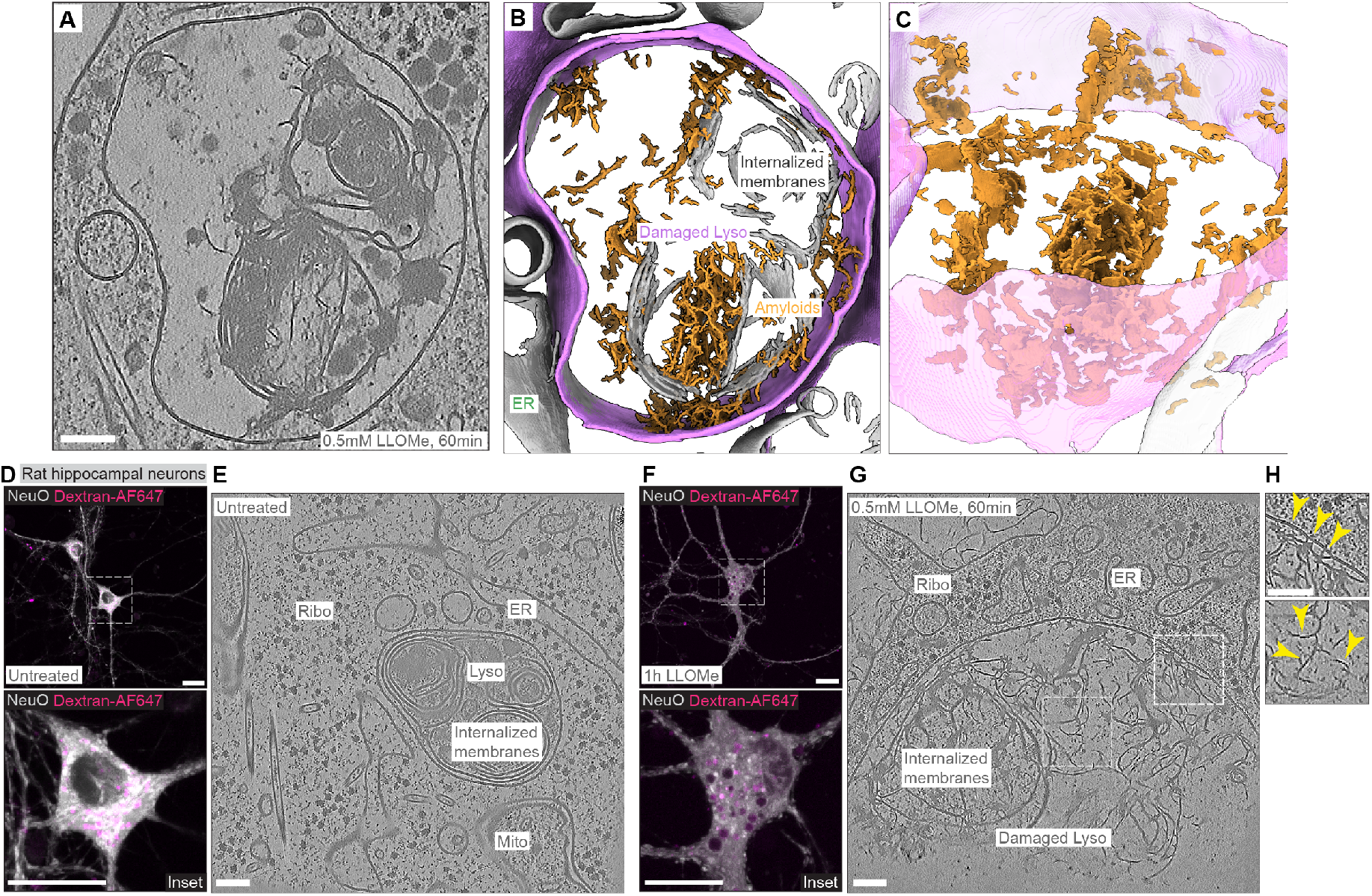
Ultrastructural analysis of LLOMe-damaged lysosomes reveals distinct electron dense fibrils inside lysosomes. (**A**-**B**) Tomographic slice of a LLOMe-damaged lysosome in HeLa and corresponding segmentation. The lysosomal membrane is labelled in purple, and fibrils are colored in orange. (**C**) Tilted view of the segmentation in B, showing sheet-like and fibrillar structures. (**D**) Live-cell imaging overview and zoom of untreated rat hippocampal neurons with lysosomes loaded with Dextran-AF647 (10kDa, magenta) and labeled with live-neuron dye NeuO in grey. (**E**) Tomographic slice of lysosomal structure inside the untreated neuron. Visible cellular structures are indicated. (**F**) Live-cell imaging overview and zoom of LLOMe-treated neurons with lysosomes labelled by Dextran-AF647 and live-neuron dye NeuO in grey. (**G**) Tomographic slice of lysosome after 0.5 mM LLOMe treatment for 1 h. (**H**) Exemplary zoom-in area showing fibrils that contact and deform the outer-most membrane. Visible fibrillar structures are indicated by yellow arrows. Scale bars: 100 nm (**A, E, G** and **H**), 20 μm (**D** and **F**).

### LLOMe forms Cathepsin C-dependent polymers that assemble into fibrillar amyloids

Having visualized the fibrillar structures *in situ*, we next sought to understand the molecular mechanisms underlying their formation. We established several *in vitro* reconstitution systems to examine LLOMe-induced membrane permeability and polymer assembly.

Previous studies have shown that CatC switches from peptidase to transferase activity upon increased pH, leading to the formation of longer peptide chains (*36–38*). We reconstituted LLOMe polymers with active CatC at pH 7.5 and used MALDI-MS to determine the minimal repeat units. Larger polypeptide chains including leucyl-leucine (LL) dimers and trimers were detected at pH 7.5, consistent with previous results (*19*) (Fig. 3A, Fig. S4A). Monitoring reaction products over time, we observed that (LL)_*2*_OMe-dimer was produced first and then consumed forming (LL)_*3*_OMetrimers as the main reaction product (Fig. 3B).

**Fig. 3.**
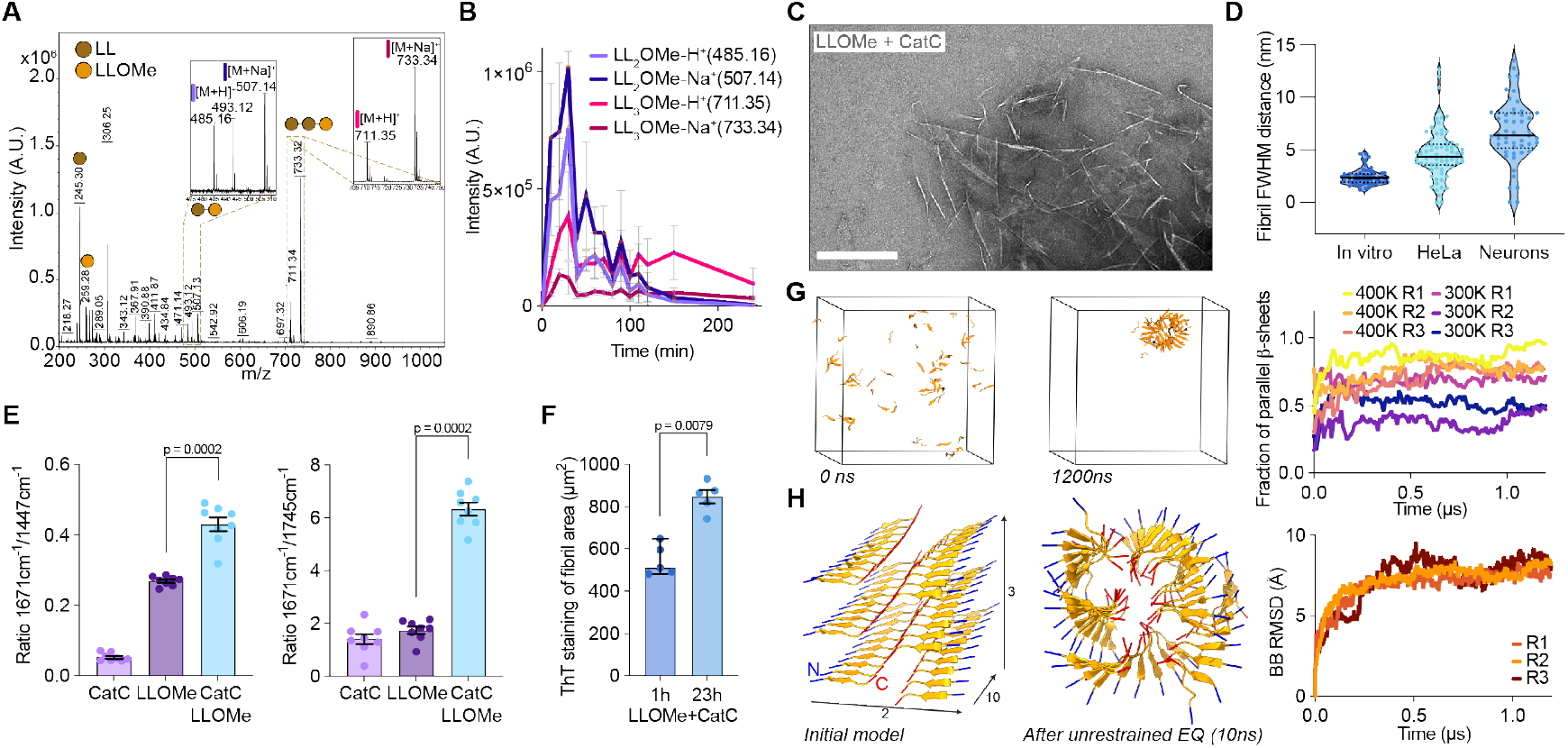
LLOMe polymerizes and forms amyloids dependent on Cathepsin C and neutral pH. (**A**) MALDI-MS spectra of LLOMe (10 mM) and active CatC (10 ng/μL) reaction mixture. Main peaks were identified as (LL)_3_-OMe and to lesser extend (LL)_2_-OMe and LLOMe. (**B**) Time-resolved MALDI-MS spectra showing LLOMe polymer intermediates over time (mean ± SD). (**C**) Negative stain EM micrograph of LLOMe (5 mM) and active CatC (5 ng/μL) reaction mixture showing amyloid fibrillar structures (scale bar: 100 nm). (**D**) Quantification of fibril full width at half maximum (FWHM) distances across negative stain EM and in situ tomograms in HeLa and neurons. Median and interquartile are displayed. (**E**) Ratios of Raman intensities at 1671cm^-1^/1447cm^-1^ and 1671cm^-1^/1745cm^-1^ between CatC (1 ng/µL), LLOMe (1 mM), and combination of both. Mann-Whitney test was conducted with p-values indicated (mean ± s.e.m., n=8). (**F**) Area quantification of ThT staining of LLOMe fibrils (n=5, mean ± s.e.m.) (**G**) MD simulation of 60 (LL)_3_-OMe that spontaneously form mixed (at 300K) or predominantly parallel (at 400K) β-sheets over three repeats. (**H**) MD simulation of assembled parallel β-sheets that spontaneously form a fibril and RMSD plot to visualize ensemble stability over three repeats. N-(blue) and C-termini (red) of monomers are indicated.

We next monitored potential higher-order oligomeric assemblies of (LL)_3_OMe formed in our *in vitro* reconstitution using negative-stain electron microscopy (Fig. 3C, Fig. S4B). We observed fibrillar structures with a diameter of approximately 3-6 nm, consistent with the *in situ* observed fibrillar structures (Fig. 3D). To determine the molecular identity of these assemblies, we employed Raman spectroscopy to characterize the bonding in LLOMe higherorder oligomeric structures. We observed a sharp amide-I band peak at 1671 cm^−1^, a hallmark of β-sheets (*39–41*) (Fig. S4C). Comparison of the peak ratios at 1671/1447 cm^−1^ (β-sheets) and 1671/1745 cm^−1^ (amide vs. esters), confirmed assembly of (LL)_3_OMe into β-sheet amyloids exclusively in the presence of CatC (Fig. 3E). To experimentally validate amyloid formation, we stained *in vitro*-derived LLOMe products with thioflavin T (ThT), a dye that selectively binds the cross-β fold of amyloid fibrils (*42*) (Fig. 3F). In the presence of CatC at pH 7.5, dense ThT-positive structures formed with increasing fluorescence intensity over time confirming the amyloid nature of higher-order (LL)_3_OMe assemblies (Fig. S4D).

We leveraged this structural information to model the amyloid architecture using computational structural biology approaches. AlphaFold3 (*43*) predicted both parallel and antiparallel pairings for inputs of 6 to 120 (LL)_*3*_OMe molecules (Fig. S4E). To gain further insight into the fibril architecture, we turned to unbiased fibril assembly using molecular dynamics (MD) simulations. At elevated temperature (400K), sixty LL_*3*_OMe molecules placed in a simulation box formed predominantly parallel β-sheets with cross-β interactions (Fig. 3G). Repeating these simulations with pre-formed parallel trimers or dodecamers and 180 LL_*3*_OMe molecules total yielded large, helical, in some cases box-spanning fibrils (Fig S4F). Integrating the structural information from experiments and computations, we built a model of the (LL)_*3*_OMe fibrils, consisting of two sheets of stacked parallel β-sheets rotated by 180 degrees with respect to each other. In this assembly, the charged N-termini are located at the fibril surface, while the neutral C-termini are stacked against each other in the core. In further simulations, a small segment of this sheet-like fibril rapidly rearranged into a helical fibril that was then stable for 1 µs (Fig. 3H, Fig. S4G). By contrast, when exploiting periodic boundary conditions to form a continuous “infinite” sheet, the fibril remained stable in its original conformation (Fig. S4H).

### LLOMe amyloids are the main cause of lysosomal membrane rupture

Notably, LLOMe amyloids observed by *in situ* cryo-ET were often in direct contact with either the lysosomal limiting membrane or internal membranes, and appeared to deform these membranes at contact sites (Fig. 4A). To quantify these deformations, we assessed membrane curvature heterogeneity on a global level by calculating the standard deviation of curvedness per lysosome. This analysis revealed no overall difference between LLOMe-treated and untreated lysosomes (Fig. 4B). However, to assess an elevation of curvedness on a local level, we next extracted patches with curvedness ≥0.021 nm^−1^ from both conditions (Fig. 4C). This revealed that the surface area of high-curvature patches was significantly increased in highly damaged LLOMe-treated lysosomes (Fig. 4D). Furthermore, high-curvature regions were spatially associated with amyloid-membrane contact sites (Fig. 4E-F, Fig. S5A), suggesting that amyloid deposition induces local membrane deformation.

**Fig. 4.**
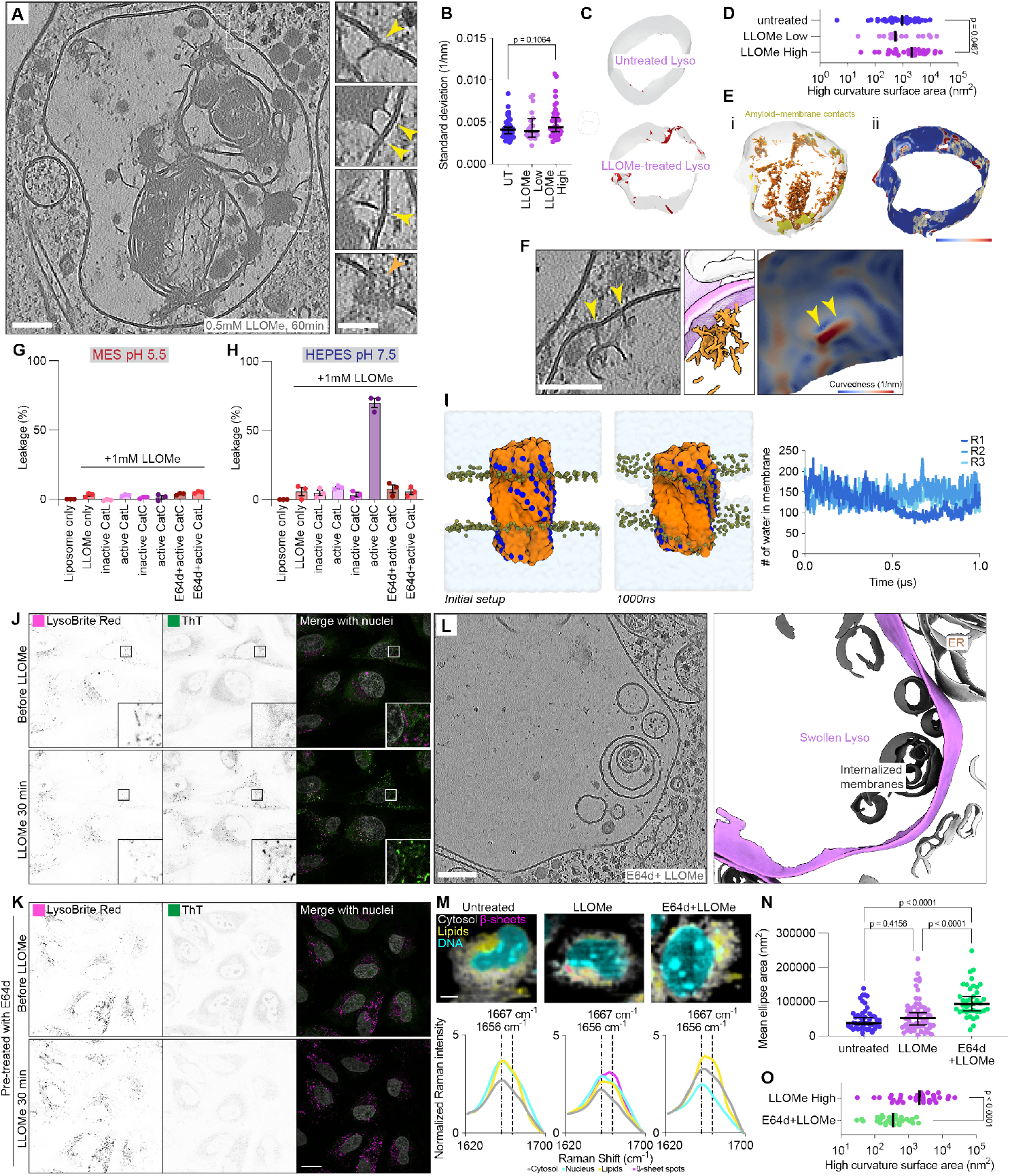
Amyloids formed after LLOMe-treatment lead to lysosomal membrane rupture. (**A**) Tomographic slice showing a LLOMe-damaged lysosome. Zoom-in views of different z-slices, shown in A, highlight membrane deformations by amyloids (yellow arrows) and a potential repair site (orange arrow). (**B**) Standard deviation of global lysosome curvedness across datasets. (**C**) Visualization of high curvature patches on untreated and LLOMe-treated lysosome membranes in red. (**D**) Comparison of high curvature patch surface areas. (**E**) Contacts of amyloids with high curvature patches. (i) Lysosome membrane patches within 15 nm of LLOMe fibrils are highlighted in yellow next to amyloid segmentation in orange. (ii) Membrane curvature measurements overlaid with the same fibril patches in yellow. (**F**) Zoom-in of tomogram with high curvature of the lysosome membrane contacting an amyloid. High curvature peaks are marked with yellow arrows. Curvedness is color-coded from 0.0008 nm-1 to 0.02 nm-1 (**E**-**F**). (**G**-**H**) Endpoint percentage of SRB leakage from LUVs (∼100nm) at pH 5.5 and pH 7.5 (n=3, mean ± s.e.m.). (**I**) MD simulation of de novo LLOMe fibril model in lysosome-like membrane forming two holes. Quantification of water molecules in inter-leaflet space over 1 µs across three repeats. (**J**) Live-cell image sequence of ThT fluorescence localizing to damaged lysosomes upon 0.5 mM LLOMe treatment, Lysosomes were labeled with LysoBrite Red. (**K**) Live-cell image sequence as in J, but with cells pre-treated with 50 μM cathepsin inhibitor E64d prior to 0.5 mM LLOMe treatment. (**L**) Tomographic slice of lysosomes from HeLa cells pretreated with E64d and treated with LLOMe (E64d+LLOMe-treated) (left) and corresponding membrane segmentation (right). (**M**) In situ Raman microscopy of untreated, LLOMe-treated, and E64d+LLOMe-treated cells with cellular components marked after Endmember analysis. Zoom-in of in situ Raman spectra between 1620 cm-1 and 1700 cm-1 showing shift of β-sheets spots towards 1667 cm-1. (**N**) Mean ellipse area of lysosomes from three different datasets (untreated n=46, LLOMe n=60, E64d+LLOMe n=39). (**O**) High curvature surface changes between E64d+LLOMe-treated and LLOMe-treated lysosomes. Scale: 100 nm (**A, C, F, L**), 50 nm (zoomed-in views in **A**), 20 μm (**J** and **K**), 5 µm (**M**). Median and 95% CI are drawn (**B, D, N**). Statistical analysis: Mann-Whitney test with p-values indicated (**B, D**); one-way ANOVA with multiple comparisons test and p-values indicated (**N**).

To elucidate the effects of LLOMe amyloids on membrane integrity, we developed a liposome leakage assay based on established protocols for small molecules and α-synuclein amyloids (*44, 45*). Sulforhodamine B (SRB) was encapsulated at self-quenching concentrations into large unilamellar vesicles (LUVs). Upon membrane disruption, SRB dilution into the external solution relieves self-quenching, resulting in an immediate fluorescence increase proportional to leakage (Fig. S5B). We investigated the membrane-disrupting capability of LLOMe in the presence of active CatC and its activator Cathepsin L (CatL) at physiological lysosomal pH (5.5). Similar to the LLOMe-only control, none of these conditions induced membrane leakage (Fig. 4G, Fig. S5C). However, when reactions were performed at pH 7.5, active CatC plus LLOMe induced a strong increase in SRB fluorescence, achieving 75% leakage relative to a detergent control (Fig. 4H). Importantly, this leakage was completely abolished by adding the CatC-inhibitor E64d (*46*) to the activated CatC (Fig. 4H), confirming that CatC-dependent LLOMe-derived amyloid fibrils can directly promote membrane rupture.

To understand the molecular mechanism of amyloid-induced membrane damage, we performed MD simulations using our *de novo*-generated helical fibril structure. We observed spontaneous insertion of the fibril into the surface of a lysosome-like membrane (Fig. S5D-E). When the fibril was positioned within the membrane, holes on both sides formed rapidly in close proximity to the fibril. The fibril disrupted the hydrophobic acyl chains of the lipid bilayer through its charged, exposed N-termini. These pores persisted throughout the complete 1 µs simulation (Fig. 4I), suggesting that once inserted into the membrane, bilayer integrity is compromised through charge interactions, leading to hole formation. Considering the osmotic pressure within lysosomes, such holes would likely expand over time if not sealed, potentially generating the large ruptures observed by *in situ* cryo-ET (Fig. 1Q).

The findings from *in vitro* reconstitution predict that CatC inhibition should block LLOMe amyloid formation and consequently prevent membrane rapture. To test this prediction, we examined the effects of E64d pretreatment on LLOMe-induced amyloid formation. Using live-cell microscopy, we monitored ThT-positive puncta in lysosomes following 30 minutes of LLOMe treatment (Fig. 4J). ThT puncta were abolished when cells were pretreated with E64d prior to LLOMe exposure (Fig. 4K), confirming CatC-dependent amyloid formation in intact cells. This finding is consistent with recent data demonstrating that the specific CatC inhibitor AZD5248 prevents amyloid formation (*47*). The protective effect of CatC inhibition was further confirmed by immunofluorescence staining against GAL-3 and CHMP2A, which showed no puncta formation upon LLOMe treatment, indicating an absence of membrane damage (Fig. S6A-B). Notably, even the combination of E64d with BAPTA-AM pretreatment, which before exacerbated LLOMe-induced damage (Fig. 1O), did not result in GAL-3 puncta formation or ESCRT recruitment to lysosomes (Fig. S6C-D), demonstrating that CatC-dependent amyloid formation is a prerequisite for LLOMe-mediated membrane damage. *In situ* Raman microscopy of cells confirmed LLOMe-dependent formation of β-sheet structures within lysosomal compartments; these signatures were absent in both untreated and E64d-pretreated cells (Fig. 4M, Fig. S7A). Finally, we investigated the ultrastructure of E64d-pretreated lysosomes after LLOMe incubation using *in situ* cryo-ET (Fig. 4L). Strikingly, we did not observe amyloid structures within these lysosomes (Fig. S7B-C); however, we noted apparent lysosomal swelling (Fig. S7B). Quantification of mean lysosomal ellipse area, perimeter and equivalent diameter revealed a significant, progressive increase from untreated to LLOMe-treated to E64d+LLOMe-treated lysosomes (Fig. 4N, Fig. S7D-E). Morphometric analysis revealed that E64d pretreatment rescued LLOMe-induced membrane alterations. Both membrane thickness and overall curvedness were reduced compared to LLOMe treatment alone, approaching values observed in untreated lysosomes (Fig. S7F-G). Furthermore, high-curvature patch analysis showed a significant decrease in the surface area of high-curvature regions in E64d+LLOMe-treated lysosomes (Fig. 4O, Fig. S7H). These data indicate that CatC inhibition prevents amyloid-mediated localized membrane deformation while the general lysosomal swelling, likely an osmotic response to LLOMe accumulation, occurs independently of amyloid formation.

Together, these results lead to a model in which CatC-dependent LLOMe amyloids physically deform and rupture the lysosomal membrane through mechanical stress, a mechanism that operates in both immortalized cell lines and primary neurons (Fig. 5).

**Fig. 5.**
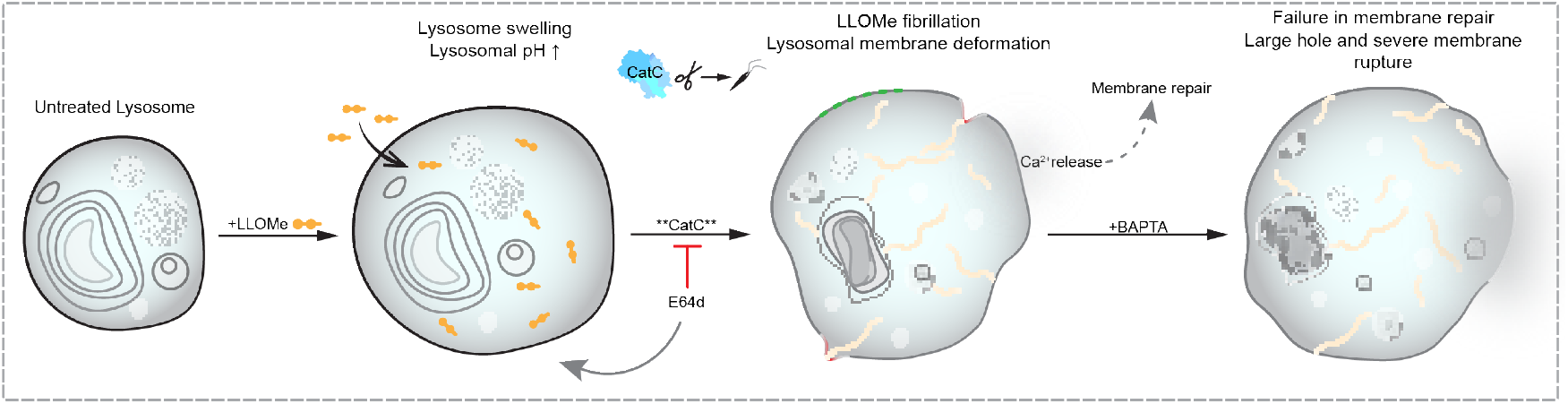
Cathepsin-mediated LLOMe amyloid formation drives mechanical rupture of lysosomal membranes. Proposed mechanism of LLOMe-induced lysosomal rupture. Cathepsin C-dependent polymerization of LLOMe generates intralysosomal amyloid structures that mechanically deform and rupture the limiting membrane. This damage is counteracted by membrane repair pathways.

## Discussion

Using cryo-ET, we have directly visualized the structural basis of LLOMe-induced lysosomal membrane damage and revealed an unexpected mechanism: CatC-dependent amyloid formation that mechanically ruptures the lysosomal membrane. This challenges the prevailing view that LLOMe damages lysosomes through simple polymerization into membrane-permeabilizing peptide chains (*15, 19*). Instead, our data demonstrate that LLOMe polymerization products assemble into sheet-like amyloid structures that physically deform membranes, creating localized high-curvature patches and ultimately generating holes ranging from 50 to 400 nm in diameter. The CatC dependence of this mechanism, confirmed through E64d inhibition both *in vitro* and in cells, establishes a new paradigm for understanding lysosomotropic agent-induced membrane damage.

Our findings reveal a dynamic interplay between damage and repair. ER recruitment scales with damage severity, and membrane repair machinery can seal small ruptures under normal conditions, explaining why catastrophic membrane failure occurs in only a small fraction of LLOMe-treated lysosomes. However, continuous LLOMe exposure or impairment of Ca^2+^-dependent repair pathways, as demonstrated by BAPTA-AM treatment, shifts this balance toward persistent membrane rupture. This suggests that the cell’s capacity to manage lysosomal damage depends critically on the rate of damage accumulation relative to repair capacity.

The amyloid-mediated mechanism we describe has important implications beyond LLOMe biology. In neurodegenerative diseases and LSDs, protein aggregates accumulate within lysosomes and are thought to compromise lysosomal function (*2*).Our demonstration that amyloid structures can directly rupture lysosomal membranes through mechanical stress provides a structural framework for understanding how disease-relevant aggregates – including α-synuclein, tau, and others – may compromise lysosomal integrity. Importantly, CatC’s transferase activity at neutral pH could be broader than currently anticipated. It is possible that distinct dipeptides generated during protein degradation within lysosomes could similarly undergo CatC-dependent polymerization and amyloid assembly if pH dysregulation occurs. These dipeptide-derived amyloids could act as seeds for further aggregation similarly to other disease-relevant amyloid fibrils (*47–49*), potentially initiating a feed-forward cycle wherein membrane damage leads to pH neutralization, promoting additional CatC-dependent amyloid formation and further membrane rupture. This raises the possibility that CatC-mediated amyloid formation from certain endogenous dipeptides could contribute to lysosomal dysfunction in diseases characterized by impaired lysosomal acidification. Moreover, the fact that CatC inhibition completely prevents LLOMe-induced damage while still allowing lysosomal swelling indicates that membrane rupture and volume changes are mechanistically separable events, with amyloid formation being the critical determinant of membrane failure. The fibrillar architecture we observe in LLOMe amyloids resembles structures formed by pathological protein aggregates (*50*), suggesting a conserved mechanism of membrane disruption.

These findings establish a new model for lysosomal membrane disruption and suggest potential therapeutic strategies. If pathological protein aggregates damage lysosomes through similar mechanical mechanisms, interventions that prevent aggregate-membrane interactions or enhance membrane repair capacity could protect lysosomal integrity in disease. Future studies examining whether disease-relevant amyloids induce similar membrane deformations, and whether their effects can be mitigated through targeted interventions, will be essential for translating these mechanistic insights into therapeutic approaches for neurodegenerative diseases and LSDs.

## Acknowledgements

We thank all members of the Mechanism of Cellular Quality Control research group, Mark Lindner, Susann Kaltwasser, Simone Prinz, and Maarten Tuijtel from the Central Electron Microscopy Facility at Max Planck Institute of Biophysics for support and advice. We acknowledge Stephan Junek and Lorenzo Latino from the Imaging Facility of the Max Planck Institute for Brain Research and Max Planck Institute of Biophysics for expertise and support. We thank the Max Planck Computing and Data Facility for providing infrastructure.

M.M and D.A.G acknowledge Hamidreza Rahmani for critical input on scripting and computational methods. We thank Jean-Christophe Ducom and Lisa Dong at The Scripps Research Institute for computational support. We thank Jürgen Köfinger for insightful discussions and feedback on the simulation setups.

We also acknowledge the use of the large language models GPT-4o/GPT 5.2 (OpenAI), Gemini 2.5 Pro (Google), and Claude Opus 4.5 (Antrophic) for writing initial code snippets and debugging errors.

The work was supported by the Max Planck Society, the European Union. Views and opinions expressed are, however, those of the author(s) only and do not necessarily reflect those of the European Union or the ERC Executive Agency. Neither the European Union nor the granting authority can be held responsible for them.

Michael J. Fox Foundation administers the grants ASAP-000350, ASAP-000282 and 024268 on behalf of ASAP and itself. For the purpose of open access, the author has applied for a CC-BY public copyright license to the Author Accepted Manuscript (AAM) version arising from this submission. This paper was typeset with the bioRxiv word template by @Chrelli: http://www.github.com/chrelli/bioRxiv-word-template

## Funding

Boehringer Ingelheim Fonds PhD Fellowship (D.L.)

German Research Foundation Research Training Group iMOL GRK 2566/1 (D.L., J.B., M.W., F.W.)

Aligning Science Across Parkinson’s (ASAP-000350, ASAP-000282/ASAP-024268) through the Michael J. Fox Foundation for Parkinson’s Research (D.L., W.Z., J.L., J.W.H., G.H., F.W.)

Pew Scholars Program (D.G.)

National Institutes of Health (NIH) grant RF1NS125674 (D.G.) and R01NS110395 (J.W.H.)

Fred and Joan Goldberg Post-doctoral Fellowship (F.K.)

ERC, IntrinsicReceptors, 101041982 (F.W.)

## Author contributions

Conceptualization: D.L., W.Z., F.W.

Methodology: D.L., W.Z., M.M., J.S., A.S., J.B., J.Br., F.K., J.P., S.W., J.La.

Investigation: D.L., W.Z., M.M., J.S., A.S., J.B., J.Br., F.K., S.O., J.Li., J.P., J.G., L.S., D.H., N.W.

Visualization: D.L., W.Z., M.M., J.S., J.Br., F.K., F.W.

Funding acquisition: D.L., M.K., J.W.H., E.S., G.H., D.G., F.W.

Supervision: S.W., J.La., M.K., J.W.H., E.S., G.H., D.G., F.W.

Writing – original draft: D.L., W.Z., F.W.

Writing – review & editing: D.L., W.Z., M.M., J.S., A.S., J.B., J.Br., F.K., S.O., J.Li., J.P., J.G., L.S., D.H., N.W., S.W., J.La., M.K., J.W.H., E.S., G.H., D.G., F.W.

## Data availability

The data, code, protocols, and key lab materials used and generated in this study are listed in a Key Resource Table alongside their persistent identifiers, which has been deposited at http://Zenodo.com under http://doi.org/10.5281/zenodo.18230779. The cryo-ET map of the flotillin complex can be accessed through EMDB-56238. All tomograms shown in the main figures are publicly available under the following accession codes: EMDB-56295 (untreated, Fig. 1A), EMDB-56296 (LLOMe-treated, Fig. 1H), EMDB-56297 (LLOMe-treated, Fig. 1P), EMDB-56298 (BAPTA AM pre-treated, LLOMe-treated, Fig. 1T), EMDB-56300(LLOMe-treated, Fig. 2A), EMDB-566327 (neuronal untreated, Fig. 2E), EMDB-56330 (neuronal LLOMe-treated, Fig. 2G), EMDB-566329 (E64d pre-treated, LLOMe-treated, Fig. 4L). The mass spectrometry data have been deposited to the ProteomeXchange Consortium (http://proteomecentral.proteomexchange.org) via the PRIDE partner repository (*51*). Anonymous reviewer access is available upon request.

## Materials and Methods

### Materials

The data, code, protocols, and key lab materials used and generated in this study are listed in a Key Resource Table alongside persistent identifiers at 10.5281/zenodo.18230779.

#### Cell culture and chemical treatment

HeLa TMEM192-3xHA (RRID: CVCL_D1KR) cells were cultured in a humidified incubator at 37℃ in 5% CO_2_ in full medium (Dulbecco’s modified Eagle’s medium supplemented with 10% fetal calf serum and 4 mM L-glutamine). Compounds were used at the following concentrations: 0.5 mM LLOMe (Cayman Chemical, CAY16008), 50 µM BAPTA-AM (ThermoFisher Scientific, B6769), E64d (Adooq, A13259) at 50 µM or live cells or 100 µM for fixed cells. Concentrated stock solutions were prepared in DMSO and stored at -20℃.

### Immunofluorescence

Cells were grown on coverslips. After the indicated treatments, fixed cell samples were prepared at room temperature. Cells were washed twice with PBS and fixed with 3% methanol-free paraformaldehyde (PFA) in PBS for 20 min, followed by permeabilization with 0.1% saponin for 10 min. Coverslips were then blocked with 5% BSA in PBS for 30 min and incubated with primary antibodies in 1% BSA in PBS for 1hr. After washing with 1% BSA in PBS, coverslips were incubated with secondary antibodies in 1% BSA for 1hr. Finally, coverslips were washed with PBS and water and mounted in mowiol.

Antibodies used for immunofluorescence are listed as follows. Primary antibodies: GAL-3 Alexa 488 conjugated (1:250, BioLegend, 125410), CHMP2A (1:500, ProteinTech, 10477-1-AP), LAMP1 (1:500, Santa Cruz Biotechnology, sc-20011). Secondary antibodies: anti-rabbit IgG Alexa Fluor™ 488 (1:1000, ThermoFisher, A-21206), anti-mouse IgG Alexa Fluor™ 594 (1:1000, ThermoFisher, A-21203). Nuclei were stained with 1µg/mL Hoechst 33342 (ThermoFischer, H1399).

### ThT staining of live cells

Cells were seeded on glass-bottom dishes (Greiner, 627860) and grown until 70-80% of confluency. Cells were incubated with 100 µM ThT in full medium overnight. Before imaging, cells were washed twice with PBS and labelled with LysoBrite Red (1:2500, AAT Bioquest, 22645) and Hoechst 33342 (1 µg/mL) in full medium for 15min. The excess dyes were wash twice with PBS and cells were incubated in full medium, prior to the live cells imaging.

### *In vitro* reconstitution of LLOMe fibrils

Recombinant human cathepsin C (CatC, proenzyme) was obtained from Acro Biosystems (CAC-H52H3) and human cathepsin L (CatL, proenzyme) was obtained from Sino Biological (10486-H08H). To activate CatC, 100 ng/µL CatC mixed with 200 ng/µL CatL was incubated in activation buffer containing 25 mM MES pH 5.5 and 2.5 mM TCEP at room temperature for 2 h. To generate LLOMe fibrils, LLOMe was dissolved in buffer containing 20 mM HEPES pH7.5 and 140 mM NaCl to a final concentration of 34 mM. All the reaction were prepared in the same buffer unless otherwise indicated.

To prepare samples for negative staining, 5 ng/µL active CatC was mixed with 5 mM LLOMe and incubated at room temperature for 2 h. For ThT staining of LLOMe polymers, 1 ng/µL active CatC was mixed with 5mM LLOMe in a final volume of 50 µL in 384-well glass-bottom plate (Greiner, 781892) and incubated for 30 min. ThT dye was then added to a final concentration of 1mM and mixed. To prevent evaporation, 20 µL mineral oil was layered on top prior to confocal imaging.

### Liposome leakage assay

100% POPC (Avanti Polar Lipid, 850457C) was prepared to a final concentration of 2 mM. The lipid mixture was dried under nitrogen gas and further vacuum-desiccated for at least 2 h to remove the residual solvents. The lipid film was resuspended in buffer containing 40 mM SRB, 10 mM HEPES pH7.5 and 70 mM NaCl with vortexing. The lipid solution was subjected to 5 cycles of snap freezing in liquid nitrogen and thawing on a heat block at 42℃. Liposomes were then extruded through 200 nm polycarbonate membrane at least 10 times, followed by extrusion through 100 nm membrane at least 20 times using a Mini-Extruder (Avanti Polar Lipid). Excess unencapsulated SRB dye was removed by a Sephadex® G-50 column pre-equilibrated with 20 mM HEPES pH7.5 and 140 mM NaCl.

For the leakage assay, 100 µL reactions containing 1 mM LLOMe and 100 nM SRB-containing liposomes at acidic or neutral pH were prepared in 96-well microplate (Greiner, 675076). The acidic assay buffer contained 20 mM MES pH5.5 and 140 mM NaCl, whereas the neutral assay buffer contained 20 mM HEPES and 140 mM NaCl. Active CatC was prepared as described above. CatL was mixed in activation buffer to a final concentration of 100 ng/µL and incubated at room temperature for 2 h to allow self-activation. Final enzyme concentrations in the assay were 1 ng/µL for CatC and 2 ng/µL CatL. E64d was added to the reaction mix after enzyme activation to a final concentration of 100µM.

All the conditions were prepared using the same batch of liposomes and measured with TECAN Spark® Microplate Reader. The SRB fluorescence (ex/em 540nm/586nm) was monitored for 2 h at room temperature. 0.4% Triton X-100 was used to define the maximum SRB leakage (100%) from liposomes, while SRB containing liposomes alone serves as a no-leakage control. The percentage of leakage was calculated as follows:

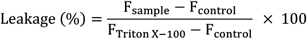

The experiments were performed with three independent repeats.

### Confocal microscopy and image quantification

All cell images were acquired using Zeiss LSM980 equipped with 63x/ 1.4NA oil immersion objective. Random fields of view were picked for imaging. For cell experiments, Hoechst 33342 was excited with 405nm laser, Alexa Fluor 488 and ThT were excited with 488nm laser, and Alexa Fluor 594 and LysoBrite Red were excited with 561nm. For fixed samples, 5 z-stacks with a step size of 0.21 µm were acquired using the same settings for each antibody staining. Live cell imaging was performed with a prewarmed imaging stage at 37℃ supplied with 5% CO_2_. ThT stained LLOMe polymers was imaged with Zeiss LSM980 with 40x/ 1.2NA water objective at room temperature. The images were acquired using excitation laser at 405nm and emission filter at 455-552nm.

All the confocal images were processed with FIJI/ImageJ (*52*). For colocalization measurement, Mander’s colocalization coefficient was calculated using JACoP plugin (https://imagej.net/plugins/jacop) (*53*) on the maximum intensity projections. 12-16 fields per condition were used for colocalization analysis. To analyze the GAL-3 puncta per cell, first a mask of the cells was created manually. Secondly, the analysis of the puncta was done in Fiji, by making maximum z-projections of the images, setting the same threshold for all of them and converting them to a binary mask. To exclude random noise only particles larger than 15 pixels were considered puncta. The counting of the puncta per cell mask was done with a python script. At least 20 cells per condition were used for spot quantification.

Areas of ThT-stained LLOMe polymers (ROIs) were identified using threshold. Integrated densities of ROIs were measured and normalized to the “LLOMe+CatC 1hr” condition. 5 fields per condition were analyzed for area and integrated density analysis.

### Mass spectrometry of *in vitro* reconstituted LLOMe fibrils

100 ng/µL active Cat C was prepared freshly as described above. For the final reaction, 10 ng/µL active CatC was mixed with 10 mM LLOMe in buffer containing 20 mM HEPES pH7.5 and 140 mM NaCl.

For each time point, to stop the reaction, 1 µl reaction setup was mixed with 1 µl matrix (2,5-dihydroxybenzoic acid (DHB), 30 mg/mL in TA50, Bruker Daltonics, Bremen) and spotted directly onto a big anchorchip MALDI target in quadruplicates. MALDI mass spectra were acquired on a Bruker Rapiflex MALDI-TOF/TOF mass spectrometer using the FlexControl 4.0 (Build 35) control software, in the mass range of 60–1500 Da in positive ion reflector mode using optimized ionization, ion optics and detector settings. For each spot, 500k shots were collected using “random complete sample” mode. Spectra were recalibrated using the near-neighbour method with a calibrant mixture (Bruker Protein Calibration standard 1, Bruker Daltonics, Bremen supplemented with an amino acid mixture (Sigma Aldrich, Germany, A6407). Full method details are available with the raw data at the PRIDE partner repository.

All spectra were evaluated using the software Bruker FlexAnalysis 4.0 (Build 14). After back-ground subtraction (TopHat), smoothing (Savitzky-Golay, width 0.05 m/z, 2 cycles) and peak picking (Centroid, s/n 3) the intensities of the (LL)_n_OMe proton and sodium adducts were extracted to monitor product formation over time. All values were calculated from 2-3 experimental and technical MALDI measurement duplicates.

### Raman microscopy

#### Drop-coating deposition Raman spectroscopy

1 mM LLOMe, 1 ng/µL CatC, and a combination of both were placed in 10 µL droplets onto calcium fluoride slides (Korth Kristalle GmbH, Altenholz, Germany) and dried for 60 min in a fume hood prior to measurement. Raman spectra were acquired using a confocal Raman microscope (alpha300R, WITec GmbH, Ulm, Germany) equipped with a 50× objective (numerical aperture 0.8, Carl Zeiss Microscopy GmbH, Jena, Germany) and a 532 nm excitation laser set to a power of 20 mW in front of the objective; the integration time was set to 4 s. For each sample, eight spectra were collected. Background subtraction and cosmic-ray removal were performed using ProjectFour software (WITec GmbH). Preprocessed spectra were then imported into MATLAB (The MathWorks Inc., Natick, United States) and subjected to further analysis. Spectra acquired from single-component analyses (LLOMe and CatC, respectively) were used as references for comparison with the mixed sample. Peak intensity ratios at 1671/1447 cm^-1^ and 1671/1745 cm^-1^ were calculated for comparative evaluation.

#### Confocal Raman microscopy of cells

HeLa TMEM192-3xHA cells were seeded on glass coverslips and incubated overnight. Cells were treated with 0.5 mM LLOMe, either directly or pre-treated with 100 µM of the cathepsin inhibitor E64d for 30 min prior to treatment with 0.5 mM LLOMe; untreated cells served as controls. After 1 h, cells were fixed with 3% PFA in PBS for 20min at room temperature and washed twice with PBS. Raman spectra were acquired using a confocal Raman microscope (alpha300R, WITec GmbH, Ulm, Germany) equipped with a 532 nm excitation laser set to a power of 40 mW in front of the objective and a 63× water-dipping objective (numerical aperture 1.0, Carl Zeiss Microscopy GmbH, Jena, Germany), yielding a spatial resolution of 333 nm. Raman scans were recorded with an integration time of 0.7 s per pixel. Background subtraction and cosmic-ray removal were performed using ProjectFour software (WITec GmbH), after which spectra were imported into MATLAB (The MathWorks Inc., Natick, United States) for smoothing (Savitzky–Golay filter, window size 9, order 3) and normalization by z-scoring. Processed data were analyzed using vertex component analysis (VCA) for endmember extraction. Endmember spectra were used to calculate abundance maps, which were overlaid in Fiji and then used for subsequent band-specific spectral analysis. Line-wise measurement artifacts were reduced by applying a Gaussian blur (σ = 1 pixel) in Fiji.

### Molecular dynamics (MD) simulation

#### AlphaFold3 modeling

We used a local AlphaFold3 (*43*) installation to predict structures of assemblies of 6, 12, 24, 36, 60, 120, 240, and 360 (LL)_3_OMe molecules. To model (LL)_3_OMe molecules, we used L_6_ as peptide input together with a CH_3_ (CCD code) ligand that was bonded to a C-terminal oxygen atom. We performed the calculation of features only once, and used them for all runs. All inferences were run with 10 seeds and 5 diffusion samples per seed.

#### Simulations of fibril formation from monomers, trimers, and dodecamers

We extracted structures of monomeric (LL)_3_OMe in β-sheet conformation, and trimeric and dodecameric parallel (LL)_3_OMe β-sheets from our AlphaFold3 predictions. We built three different systems: i) 60 monomers, ii) 60 trimers (180 peptides total) or iii) 15 dodecamers (180 peptides total) were randomly placed in a cubic box with an edge length of ∼15 nm. We solvated the systems in TIP3P water and 150 mM NaCl. We used the charmm36m force field (*54*) and TIP3P water with modified hydrogen ε parameters (*54, 55*) for all simulations of non-membrane systems. We set charged N-termini and methylated C-termini. We used Gromacs (version 2023.2) (*56*) as MD engine.

Before production, we performed one minimization and six equilibration steps, of which the first was in the NVT and all consecutive ones were in the NPT ensemble. For minimization, we used the steepest descent algorithm until the maximum force acting on any atom in the system was below 1000 kJ mol^-1^ nm^-1^. For all equilibration and production steps, we used a leap-frog integrator. For NVT equilibration, we used a Berendsen thermostat (*57*) with a target temperature of T = 300K (monomers only)/400K (monomers, trimers, dodecamers) and a characteristic time τ_T_ = 0.1 ps. For the first NPT step, we increased the characteristic time of the thermostat to 1.0 ps, and added pressure coupling via an isotropic Berendsen barostat (*57*) with a target pressure p = 1 bar, a compressibility of 4.5 · 10^-5^ bar^-1^, and a characteristic time of τ_p_ = 5 ps. For all following equilibration steps, we used a v-rescale thermostat (*58*) and a Parrinello-Rahman barostat (*59*). We used an integration time step of 1 fs for the first three equilibration steps and a time step of 2 fs for the last three. We ran the equilibrations for 1 ns, 1 ns, 2 ns, 4 ns, 4 ns, and 10 ns, respectively. We applied position restrains on heavy atoms and gradually released them over the course of equilibration. Starting from a force constant of 1000 kJ mol^-1^ nm^-2^ on backbone (including the C-terminal oxygens and the carbon atom of the methyl capping group) and 500 kJ mol^-1^ nm^-2^ on sidechain atoms during step 1 to no restraints in the last step.

We ran production simulations at the same settings as the last equilibration step for ∼1.2 µs (monomers) / ∼1.1 µs (trimers) / ∼1.0 µs (dodecamers). All simulations used a Verlet cutoff scheme (*60*), a cutoff of 1.2 nm with a force-switch modifier at 1.0 nm for Van-der-Waals interactions, a cutoff of 1.2 nm for Coulomb interactions, and Particle Mesh Ewald (*61*) for long-range electrostatics. To describe bonds with hydrogens we used the LINCS algorithm (*62*). We ran all systems in triplicates starting from different original random placements of monomers/trimers/dodecamers in the simulation box.

#### Simulations of fibril sheet fragments

We modeled the fibril sheet in ChimeraX (version 1.6.1) (*63*). We aimed for a model with charged N-termini at the outside, parallel β-sheets, a width of 4 to 5 nm, the potential for unlimited growth in length and height, and the ability to twist to minimize the exposed hydrophobic surface. We built two models with the respective dimensions 2 (width) · 3 (height) · 10 (length) and 2 · 3 · 18 peptides. We performed an energy minimization step before solvating the system in TIP3P water and 150 mM NaCl. We performed MD simulations on this fibril sheet fragment with the same settings as used for the simulation of fibril formation with a temperature of T = 300K, with one change for the 2 · 3 · 18 system: Here, we ran the last equilibration step for 30 ns instead of 10 ns. We ran all systems in triplicates for ∼1.2 µs (2 · 3 · 10) / ∼0.5 µs (2 · 3 · 18) starting from different in-vacuum energy minimizations of the respective initial fibril.

#### Simulations of “infinite” fibril sheet

We modeled the “infinite” fibril sheet by first extending the original 2 · 3 · 10 assembly (after the fourth equilibration step) to a 2 · 5 · 10 assembly using VMD (*64*). For this, we aligned the top layer of one 2 · 3 · 10 assembly to the bottom layer of another one. We then aligned the height dimension of the fibril layer to the z-axis of a simulation box and set the z dimension of the box to 5 times the inter-layer distance.

We used the exact same minimization, equilibration, and production settings as in the simulations of the fibril sheet fragments with two exceptions. We used a semiisotropic c-rescale (*65*) barostat for pressure coupling, and for production runs we used a different Gromacs version (2025.2 instead of 2023.2). We ran all systems in triplicates for 1 µs starting from different in-vacuum energy minimizations of the respective initial fibril.

#### Simulations of membrane systems

We used CHARMM-GUI (*66–68*) to construct two different membrane systems: i) The equilibrated 2 · 3 · 10 assembly placed 0.5 nm above a lysosome-like (*69*) membrane containing 204 lipids in each layer and ii) the equilibrated 2 · 3 · 18 assembly placed in the membrane. In all setups, we aligned the principal axis of the fibril with the z-axis of the box. We solvated the system in TIP3P water and 150 mM NaCl. We used the charmm36m force field and unmodified TIP3P water for all simulations of membrane systems. We set charged N-termini and methylated C-termini. We used Gromacs (version 2023.2 for minimization and equilibration, and 2025.2 for production) as MD engine.

Before production, we performed one minimization and six equilibration steps, of which the first two were in the NVT and all consecutive ones were in the NPT ensemble. For minimization, we used the steepest descent algorithm until the maximum force acting on any atom in the system was below 1000 kJ mol^-1^ nm^-1^. For NVT equilibration, we used a v-rescale thermostat with a target temperature of T = 300K and a characteristic time τ_T_ = 1.0 ps. For the first NPT step, we added pressure coupling via a semiisotropic c-rescale barostat with a target pressure p = 1 bar, a compressibility of 4.5 · 10^-5^ bar^-1^, and a characteristic time of τ_p_ = 5 ps. We used an integration time step of 1 fs for the first three equilibration steps and a time step of 2 fs for the last three. We ran the equilibrations for 1.25 ns, 1.25 ns, 1.25 ns, 5 ns, 5 ns, and 5 ns, respectively. We applied position restrains on heavy atoms and gradually released them over the course of equilibration. For the first step, we used position restraints of 4000 kJ mol^-1^ nm^-2^ on backbone atoms (excluding the C-terminal oxygens and the carbon atom of the methyl capping group), 2000 kJ mol^-1^ nm^-2^ on sidechain atoms, and 1000 kJ mol^-1^ nm^-2^ on the z coordinate of lipid phosphorus atoms (oxygen for cholesterol), as well as dihedral restraints of 1000 kJ mol^-1^ rad^-2^ on selected lipid dihedral angles. For the last step, we used a restraint of 50 kJ mol^-1^ nm^-2^ on only backbone atoms. In every system, we kept the backbone heavy atoms of 6 peptides (one per sheet) at the distal side with respect to the membrane restrained at their position with a force constant of at least 1000 kJ mol^-1^ nm^-2^ during equilibration and production runs. For system i), we also performed a set of simulations without position restraints in production runs.

Production simulations used the same settings as the last equilibration step except for the position restraints, which we turned off except for selected atoms as specified in the previous paragraph. All simulation settings not described in this section were the same as for the solution systems. We ran all simulations for 1 µs in triplicates starting from different minimizations.

#### Analysis of simulations

We computed the fraction of parallel β-sheets in the simulations of fibril formation via a heuristic using H bonds of in-register parallel and antiparallel (LL)_3_OMe β-sheets. For an in-register parallel β-sheet, H bonds are formed between the N-H group of residue i (=1,2,3,…,length(monomer)) in one monomer and the C=O group of residue i-1 in another monomer. For an in-register antiparallel β-sheet, H bonds are formed between the N-H group on residue i in one monomer and the C=O group of residue length(monomer)+1-i. With this heuristic and for a monomer of 6 residues, the pairing of N-H from residue 4 and C=O from residue 3 is ambiguous, so we excluded it. This leaves 4 possible H bonds for the parallel and 5 for the antiparallel arrangement. We corrected for this imbalance by weighting the number of antiparallel H bonds with a factor of 0.8. In summary, we calculated the fraction of parallel β-sheets *f*_*p,sheet*_ as

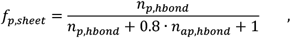

where *n*_*p,hbond*_ is the number of parallel H bonds, *n*_*ap,hbond*_ is the number of antiparallel H bonds, and 1 is added to the denominator to avoid division by 0. We used the dssp (*70*) definition for H bonds in these calculations.

We evaluated the stability of the fibril sheet structures (both fragments and “infinite” sheet) via root mean square deviation (RMSD) of backbone atoms with respect to the configuration at time t = 0. For this, we post-processed all trajectories to remove jumps across the periodic boundary of the box and keep the fibril as one unit. We then aligned the fibril structure of each frame on the reference before calculating the RMSD. In one replica (R3) of the 2 · 3 · 10 assembly, one peptide detached from the assembly, diffused across box boundaries and reattached to the fibril. This led to a poor alignment and strongly inflated RMSD values. Hence, we excluded this peptide from the RMSD analysis.

To quantify the insertion of the fibril into the membrane or, in systems where the fibril was being held in place by restraints, the recruitment of the membrane to the fibril, we computed the distance between the z coordinate of the center of mass (CoM) of the protein and z coordinate of the CoM off all phosphorus atoms. For this, we post-processed all trajectories to remove jumps of peptides across the periodic boundary of the box in z direction.

We measured pore stability in the simulations of the inserted fibril by counting the number of water molecules within the membrane every frame. We defined the number of waters in the membrane as number of water oxygen atoms with a z coordinate within 0.5 nm of the z coordinate of the CoM of all membrane phosphorus atoms.

We used 1 ns steps for every analysis, except for the calculation of *f*_*p,sheet*_, which we only did every 10 ns. We implemented the analysis in Python3 (https://www.python.org/downloads/release/python-3109/), using numpy (*71*), matplotlib (*72*), and MDAnalysis (*73*). We used VMD and ChimeraX to create visual representations of simulation systems and to build models.

### *In situ* cryo-electron tomography (cryo-ET)

#### Grid preparation

HeLa TMEM192-3xHA cells were cultured for *in situ* cryo-ET as follows: gold grids coated with holey Silicon Dioxide R1/4 at mesh 200 (Quantifoil) were glow dischared (90s, 15 mA, at 0.37 mbar), coated with 0.25 µg/mL laminin (Sigma-Aldrich, L4544) for 1h under the UV light of a sterile environment and washed twice with PBS. One day before plunge-freezing, laminin-coated grids were transferred to 8-well µ-Slide dishes (Ibidi, 80826) and ∼12,000 cells were seeded per well together with 0.1mg/mL 10kDa Dextran coupled to Alexa Fluor 647 (Dextran-AF647) (Thermo Fisher Scientific, D22914) for lysosome detection under cryogenic conditions. After overnight incubation, the cells were pre-treated with E64d (*46*) or BAPTA-AM or none prior to LLOMe (0.5 mM) incubation for 1h. Then, the grids were vitrified with PBS-diluted autofluorescent 1 µm diameter Dynabeads (Thermo Fisher Scientific, acid65011) for fluorescence correlation after 6.5 s blotting time at 37°C and 70% relative humidity in liquid ethane (at -184°C) using an EM GP2 plunger (Leica Microsystems). After plunging, the grids were clipped into autogrids with a cutout for FIB-milling.

For cryo-ET of rat neurons, grids were prepared as follows: First, electron microscopy support grids (Quantifoil™ R 2/2 or R1/4 on 200 gold mesh) were glow discharged for 90 s at 15 mA and 0.37 mBar pressure, then transferred to a sterile laminar flow hood and UV sterilized for 30 min. One grid each was placed into a glass bottom dish (Mattek, P35G-1.5-14-C) and coated with 0.1 mg/ml poly-D-lysine (BD 354210) over night. Dishes containing grids were rinsed three times with sterile ddH_2_O. The next day, rat hippocampal neurons were prepared from P0 or P1 rat pups (Sprague Dawley, IGS, Crl:CD(SD), Charles River Laboratories, RRID:RGD 734476) as described previously (*74*). First, the hippocampus was dissected, digested with Papain (Sigma, P3125), and 35,000 cells were plated in 250 µl normal growth medium (NGM; Neurobasal A without phenol red (Gibco, 12349015)) per dish. After 4 h, 500 µl glia-conditioned medium was added and cells were cultured for 21 days at 37 °C in 5% CO_2_ with the addition of 700 µl NGM supplemented with B27 (Invitrogen/Gibco, 17504-044) and GlutaMAX (Gibco, 035050-038) every week.

On DIV20, 1.5 ml medium was removed and kept in the incubator. Dextran-AF647 (0.05 mg/mL, Thermo Fisher Scientific, D22914) was added to the remaining medium and incubated overnight. On DIV21, for each grid the previously removed 1.5 ml medium was supplemented with a neuron-specific live-cell dye (NeuroFluor™ NeuO, 0.25 µM, Stemcell Technologies, 01801) (*35*), 20 mM HEPES, and LLOMe (0.5 mM). The remaining medium was replaced with the dye-supplemented medium and incubated for 20 min at 37 °C, 5 % CO2. Each grid was then imaged on a Zeiss LSM 880 Examiner upright laser scanning confocal microscope, operated at 37°C, with a 20x/0.5NA water dipping objective (Zeiss W N-Achroplan 20x/0.5 M27, 420957-9900-000). A 488 nm excitation was used and emitted fluorescence as well as transmitted light were collected as 8 x 8 tile-scan with multiple z planes at a pixel size of 415 nm x 415 nm and 3.71 µm optical sections. These confocal volumes were then stitched and maximum z-projected in the Zeiss ZEN software for subsequent correlation.

After imaging, cells were transferred in an insulated transport container on a 37°C heat pad. Grids were mounted in a Leica EM GP2 plunge-freezer maintained at 37°C 70% humidity, 2 µl of fresh medium were added to the front of the grid, and blotted from the backside for 10 s using Whatman filter paper 1. The cells were then plunged into liquid ethane kept at -184°C and stored in liquid nitrogen until further use. The total time cells were in medium containing NeuO and LLOMe was 1 hour.

#### Focused ion beam (FIB) milling

Electron-transparent lamellae were produced in a Aquilos2 dual-beam cryo-Focused Ion Beam and scanning electron microscope (cryo-FIB/SEM) instrument (Thermo Fisher Scientific). Intracellular lysosome fluorescence by Dextran-AF647 was observed using the integrated fluorescence light microscope (iFLM) of the Aquilos system. Fluorescent stacks were acquired using the 470 nm and 625 nm channels (4% laser intensity, 30ms exposure per slice, z-stack height of 6 µm, 15 slices). Correlation of the fiducial beads from the fluorescence 2D projection image (HeLa cells) or the pre-vitrification confocal fluorescence images (neurons) with the grid’s respective SEM overview image was carried out in MAPS v.3.28 (Thermo Fisher Scientific).

Semi-automatic FIB-milling was done using AutoTEM v.2.4.2 (Thermo Fisher Scientific) at a milling angle of 8° for HeLa cells and 11° for neurons. In short, the milling consisted of the following steps: i) rough milling at 1 nA to 1 µm thickness, (ii) 0.5 nA to 750 nm, and (iii) 0.3 nA to the 500 nm. Afterwards, the lamella was polished at a current of 50 pA to 150 nm) and 10 pA to 130 nm (with 0.2° overtilting). In some cases, remnants of the cell top surface with its organometallic layer had to be removed with manual intervention in addition to make the full tilt range in cryo-ET accessible.

#### Tilt series acquisition

Tilt series of HeLa cells were acquired on a Titan Krios G2 (Thermo Fisher Scientific) at 300 kV with BioQuantum energy filter and K3 direct electron detector (Gatan) at a nominal magnification of 42,000 X corresponding to a pixel size of 2.176Å/px with a 20 eV energy-slit using SerialEM v4.1–v4.2 (*75*). The tilt range was set from -52° to +68° relative to the lamella pretilt with 2° increments and a target defocus of -2.5 µm to -5 µm. The dose rate was set to ∼2.5 e/A2, resulting in ∼150 e/A2 for the complete tilt series. 10 frames per individual tilts were recorded and on-the-fly motion corrected in SerialEM.

Tilt series of rat neurons were acquired on Krios G4 at 300 kV with Selectris X energy filter and Falcon 4i camera (Thermo Fisher Scientific) using SerialEM v.4.1–v.4.2 at a nominal magnification of 42,000 X corresponding to a pixel size of 3.037Å/px with a 10 eV energy-slit using SerialEM v4.1–4.2. The tilt range was set from -49° to +69° relative to the lamella pretilt with 2° increments and a target defocus of -4 µm to -7 µm. The dose rate was set to ∼2.5 e/A2, resulting in ∼150 e/A2 for the complete tilt series. 10 frames per individual tilts were recorded and on-the-fly motion corrected in SerialEM.

The positions for tilt series acquisition were determined by visual inspection of 8,700X magnification lamella montage maps.

#### Tomogram reconstruction and denoising

The tilt series mrc stacks were corrected for dose exposure and projections of low-quality were removed manually. The tilt series was aligned using patch-tracking and reconstructed at bin4/bin6/bin8 using AreTomo2 (v.1.0.0) (*76*) and IMOD (v.4.12.62) (*77*). Tomograms were filtered using a SIRT-like filter in IMOD for display purposes.

All membrane segmentations were done using Membrain-Seg v.0.0.8 (https://github.com/teamtomo/membrain-seg) using the publicly available pretrained model (v10_alpha). Membrane segmentations of 20 tomograms of each HeLa cell dataset (untreated, LLOMe, E64d+LLOMe) were manually cleaned using Napari (*78*) to remove falsely connected membranes. All membrane segmentations were visualized using ChimeraX v.1.10 4 (*63*).

Tomogram denoising for purely visualization purposes were done at bin4 using the recently published Isonet2 method using default parameters including missing-wedge correction and CTF correction (*79*).

The tomograms were also reconstructed in Warp v.1.0.9 (*80*) using the alignments generated by AreTomo2 for easier combability with the sub-tomogram averaging workflow.

#### Template matching and sub-tomogram averaging

Template matching to identify the positions of the flotillin complex in Warp 3D-CTF-corrected tomograms was performed using GapstopTM (*81*) using a preliminary flotillin map from manual picking. After cleaning the score maps using napari_tm_particle_extraction (https://gitlab.mpcdf.mpg.de/mpibr/scic/napari_tm_particle_extraction/), the hits were extracted using Warp at bin4 (8.704Å/px). After 3D classification in Relion v.3.1.4 (*82, 83*) without symmetry, one class resembling the flotillin complex was selected and refined, re-extracted at bin1, and rerefined to generate the final map of 18.74Å resolution. For the final refinement a point symmetry of C22 was imposed, similar to the reported single-particle reconstruction (*84*).

### Lysosome morphometrics analysis

#### Surface reconstructions

20 cleaned voxel segmentations per condition (untreated, LLOMe, LLOMe+E64d) binned 8 time (17.41 Å x 17.41 Å x 17.41 Å) from Membrain-Seg v.0.0.9 (*85*) were exported and individual membranes were labeled with (Naprari plug in select-seg, https://github.com/bwmr/napari-segselect) and subsequently input into the Surface Morphometrics pipeline (*86*). Each membrane voxel segmentation was reconstructed as triangulated surface meshes using the “segmentation_to_mesh.py”. All surfaces were generated with the following parameters: maximum of 200,000 triangles, a reconstruction depth of 8, and an extrapolation distance of 1.8 nm. Edge filtering was performed on all surfaces removing all triangles within 8 nm from the edge of each surface using “distance_to_edge.py”. All downstream analyses were performed on edge filtered surfaces.

#### Curvature estimations and distance estimations

The surface morphometrics pipeline was used to generate curvature and organelle distance measurements. Pycurv (*87*) based curvature estimations of triangulated surface meshes were run using “run_pycurv.py. Curvature filtering was used for generation of high curvature patch regions within lysosomes and compared across treatment conditions. High curvature patch cut off: 0.021 nm^-1^

Inter-organelle distances were measured using “membrane_distance_orientation.py”. From resulting distance estimations, we calculated the minimum distance between a lysosome triangle surface mesh and each organelle triangle surface mesh (mitochondria, endoplasmic reticulum, and endosomes). Distance filtering was used to analyze surface area of patches of membrane in proximity to a lysosome.

#### Thickness measurements

Thickness measurements for each lysosomal surface were performed using Surface morphometrics (*88*) on SIRT filtered binned 6 tomograms (13.06 Å x 13.06 Å x 13.06 Å). Per triangle within the surface mesh a voxel density scan is performed along the normal vector from 10 nm in the positive direction to 10 nm in the negative direction using “thickness_scan.py”. From the voxel density line scan, membrane thickness is estimated by measuring the distance of the two gaussian peaks characteristic of the phospholipid head groups within the lipid bilayer. These thickness measurements are calculated by running “thickness_plots.py”.

#### Fibril associated membrane patch extraction and membrane measurements

Fibril associated patches were extracted by using Dragonfly (*89*) segmentation of LLOMe fibrils and calculating the closest distance from a fibril (all 1 values within the fibril .mrc file) to the nearest surface mesh triangle of the lysosome membrane using “surface_distance_from_mrc.py”. This distance measurement is updated in the surface mesh (vtp) as a membrane feature. This membrane feature property was then utilized for illustration, filtering and morphometrics calculations. Fibril associated patches (triangles within 5 nm of a fibril) were then extracted in ParaView (*90*) and manually inspected in ChimeraX (*63*) with the fibril segmentation to identify patches with orthogonal contact to a fibril. For per patch quantification fibril associated patch (5 nm from fibril) and proximal patch (5.1–10 nm distance from fibril)

### Lysosome volume analysis

#### Segmentation Image Processing, Skeletonization and Parametrization

Segmented lysosome membrane volumes were loaded from MRC format files. Binary segmentation masks were processed slice-by-slice along the z-axis. For each 2D cross-section, membrane boundaries were reduced to single-pixel-wide centerlines using morphological thinning to obtain skeleton representations of the membrane contours. Skeleton points were ordered along the curve by nearest-neighbor traversal starting from detected end-points (points nearest to image boundaries). Cubic B-splines were fitted to the ordered point sequences using scipy.interpolate.splprep with automatic smoothing parameter estimation based on point density. Splines were sampled at 200 uniformly-spaced parameter values to generate smooth boundary representations. Spline fits with arc lengths below 50 pixels were excluded from downstream analysis. All boundaries were treated as open arcs to accommodate partial membrane cross-sections resulting from field-of-view clipping.

#### Morphometric Measurements

For each valid 2D cross-section, we computed: (1) perimeter as the spline arc length; (2) signed curvature analytically from spline first and second derivatives using the formula κ = (x′y″ ™ y′x″) / (x′^2^ + y′^2^)^(3/2); (3) bounding box dimensions and aspect ratio; (4) centroid coordinates as the mean of spline sample points. Two-dimensional ellipses were fitted to spline sample points using a two-stage approach. Initial parameter estimates were obtained via the direct least-squares method of Fitzgibbon et al. (*91*).These estimates were refined using orthogonal distance regression (ODR) implemented via scipy.optimize.least_squares with Levenberg-Marquardt optimization, minimizing perpendicular distances from points to the ellipse boundary. Ellipse parameters comprised center coordinates (cx, cy), semi-major and semi-minor axes (a, b), and rotation angle (θ). Fit quality was assessed by the coefficient of determination (R^2^) and root mean square error (RMSE) of residual distances.

#### Bootstrap Confidence Intervals and Quality Assessment

Parameter uncertainties were estimated via bootstrap resampling with 500 iterations. For each iteration, spline sample points were resampled with replacement and ellipse fitting was repeated. The 95% confidence intervals for semi-axis lengths were computed as the 2.5th and 97.5th percentiles of the bootstrap distribution. Resampling was performed with a fixed random seed (42) to ensure reproducibility. Ellipsoid fit confidence was classified as: High (R^2^ > 0.5 and z-coverage > 70%), Medium (R^2^ > 0.3 and z-coverage > 50%), or Low otherwise. Z-coverage was defined as the fraction of the ellipsoid z-extent spanned by valid slice data. Measurements from low-confidence fits were optionally excluded from statistical comparisons.

Condition comparisons were performed using independent samples t-tests when both groups passed the Shapiro-Wilk normality test (α = 0.05), otherwise Mann-Whitney U tests were applied. Effect sizes were quantified using Cohen’s d. For multiple metric comparisons, p-values were corrected for false discovery rate using the Benjamini-Hochberg procedure.

#### Software Implementation

All image processing and statistical analyses were performed using custom Python code (version 3.11) built on NumPy (v1.24+), SciPy (v1.11+), scikitimage (v0.21+), pandas (v2.0+), joblib and Matplotlib (v3.7+).

The lysosome morphometric analysis pipeline is available as open-source software at https://gitlab.mpcdf.mpg.de/wilflinglab/lysosome-segmentation-analysis. The repository includes: Complete Python source code for all analysis steps, example datasets demonstrating input format requirements Jupyter notebook (lysosome_morphometrics.ipynb) with executable workflow, and configuration files for figure generation

### AI Assistance

The computational pipelines and analysis scripts were developed with assistance from GitHub Copilot (powered by Claude Opus 4.5), GPT-4o/GPT-5.2 (OpenAI), and Gemini 2.5 Pro. The AI tools were used for: Code drafts and optimization (subsequently verified, tested, and refined by the authors), documentation, debugging and refactoring. All AI-generated content was critically reviewed, validated against expected mathematical behavior, and tested on experimental data by the authors. The scientific methodology, experimental design, data interpretation, and conclusions are solely the responsibility of the authors.

---

**Fig. S1.**
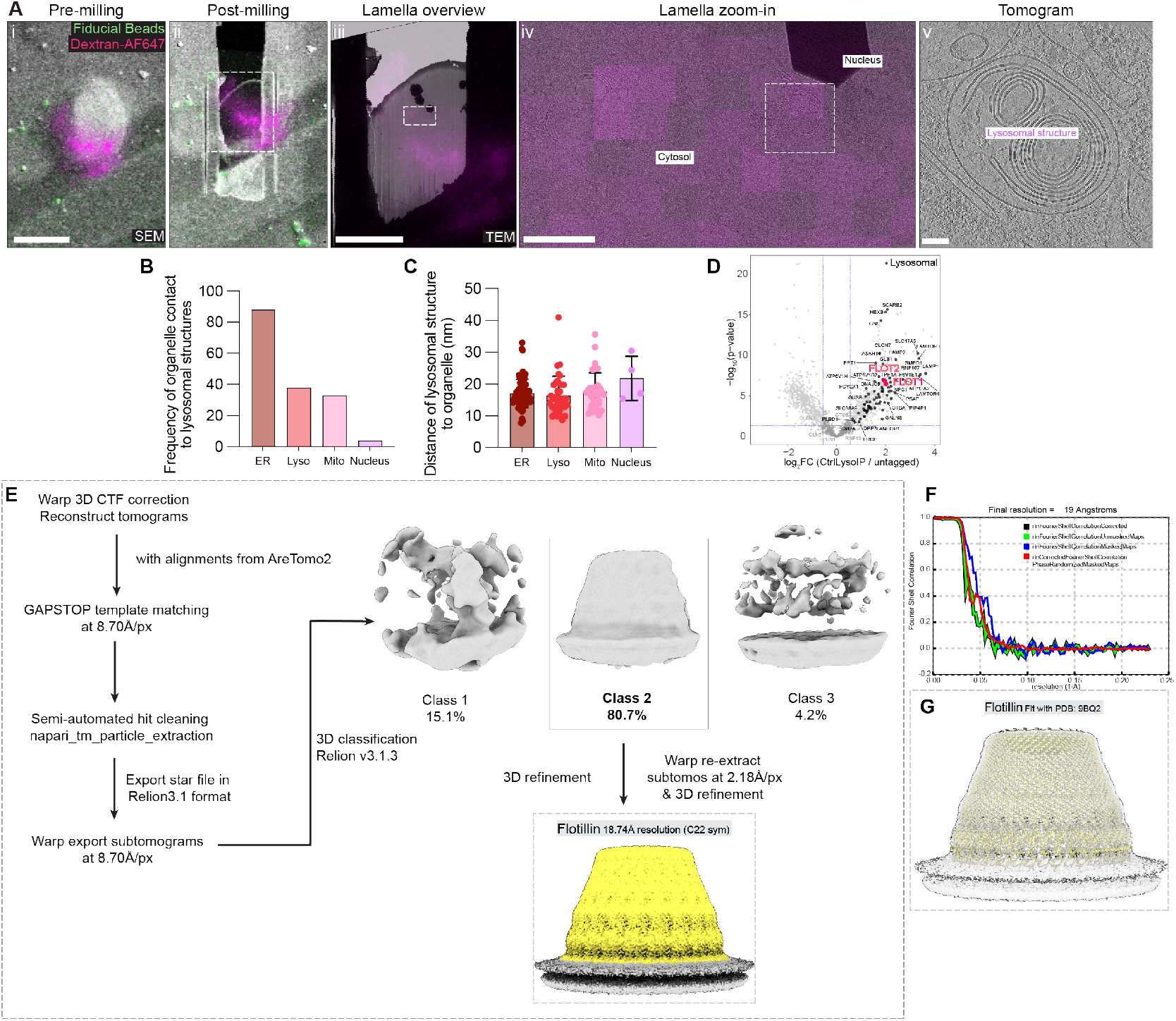
In situ investigation of lysosomes in untreated HeLa cells. (**A**) Correlative workflow to target endo-lysosomal structures in HeLa cells vitrified on cryo-EM grids. (i) maximum intensity projection overlay of Dextran-AF647 and fiducial beads fluorescence with SEM image. Scale: 20 µm (ii) Pre-milling fluorescence overlaid with final post-milling lamella SEM image. (iii) Lamella overview montage image from TEM overlaid with pre-milling fluorescence image. Distinction of cytosol/nucleus are indicated Scale: 10 µm. (iv) Zoom-in into the lamella map with tilt series acquisition area outlined. Scale: 1 µm. (**B**) Manual counting of organelle contacts with untreated lysosomes within 50 nm (**C**) Manual measurements of organelle–lysosome distances (n=65 tomograms). Median and 95% confidence interval (CI) is plotted. (**D**) Volcano plot of HeLa Lyso-IP proteomics (log_2_FC vs p-value) with lysosomal proteins marked in black and Flotillin complex (FLOT1/2) in red. (**E**) Workflow for template matching and subtomogram averaging of Flotillin complex in situ. Software versions used are indicated. (**F**) Fourier shell correlation plot from Relion output demonstrating a final resolution of 19 Å for the Flotillin cryo-ET map. (**G**) Fit of cryo-ET map (in transparent) with published atomic model of Flotillin complex PDB: 9BQ2 (*84*).

**Fig. S2.**
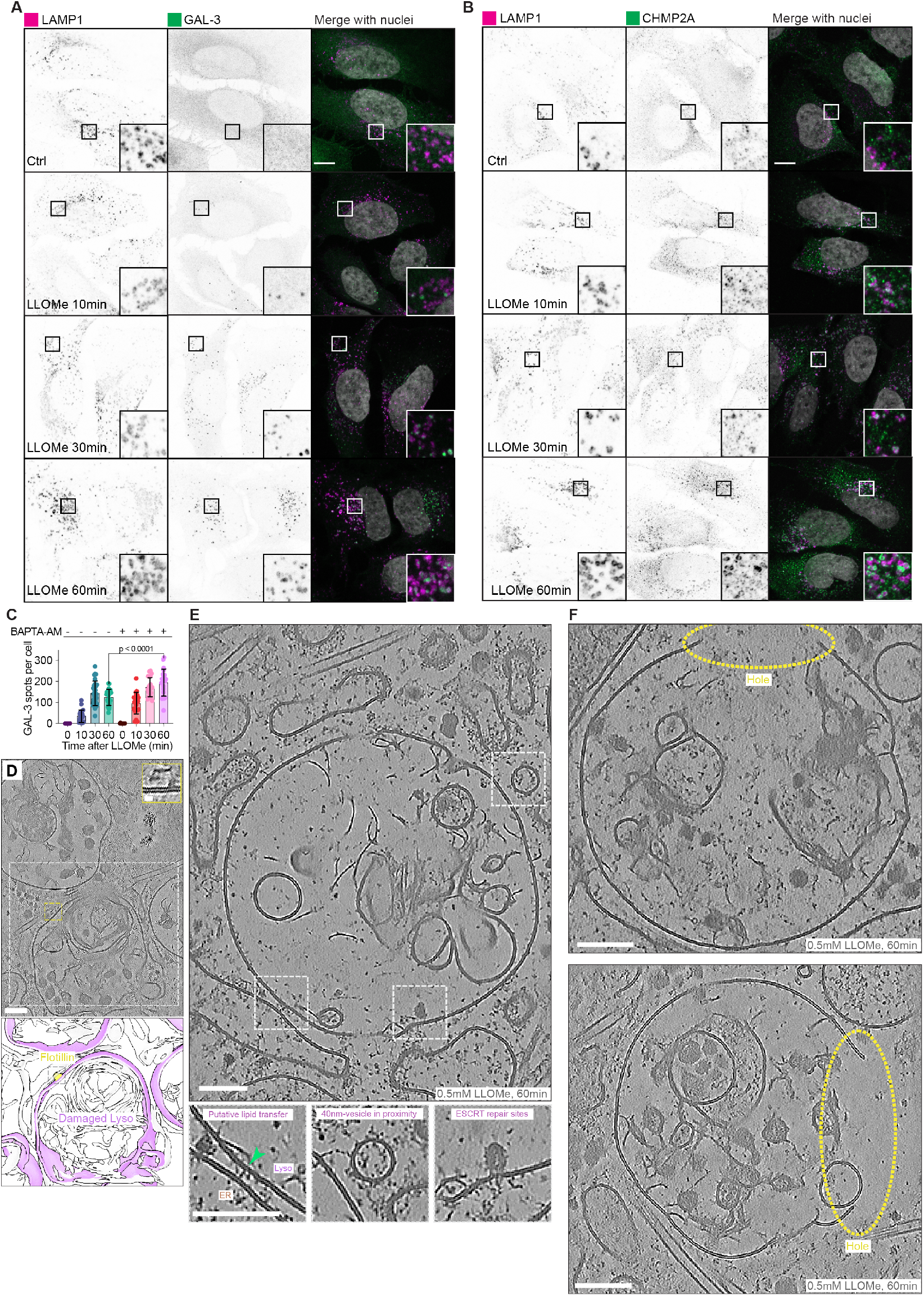
Cellular responses after LLOMe treatment. (**A**) Confocal images of immunofluorescence against LAMP1 (lysosome) and GAL-3 (lysosomal damage marker). Split channels and a merged image with Hoechst (nucleus) are shown. Untreated control and timepoints (10, 30, 60 min) after LLOMe treatment (0.5 mM) are indicated. (**B**) Same timepoints and treatments as in A but stained for CHMP2A. Scale for IF panels: 10 µm. (**C**) Quantification of GAL-3 and CHMP2A spots per cell across time course. BAPTA-AM pre-treated conditions are indicated (mean ± s.d.). Statistical test: Unpaired t-test with p-values indicated. (**D**) Tomographic slice of LLOMe-treated lysosome with segmentation and Flotillin coordinates placed back in. Zoom-in showing Flotillin in tomographic slice. Scale: 10 nm for zoom-in, 100 nm for tomographic slice. (**E**) Tomographic slice shown in Fig. 1F with zoom-ins of putative membrane repair mechanisms (lipid transfer, 40 nm ATG9A vesicles, ESCRT budding). Scale: 100 nm. (**F**) Tomographic slice of lysosomes in LLOMe-treated cells with apparent holes (circled in yellow). Scale: 100 nm.

**Fig. S3.**
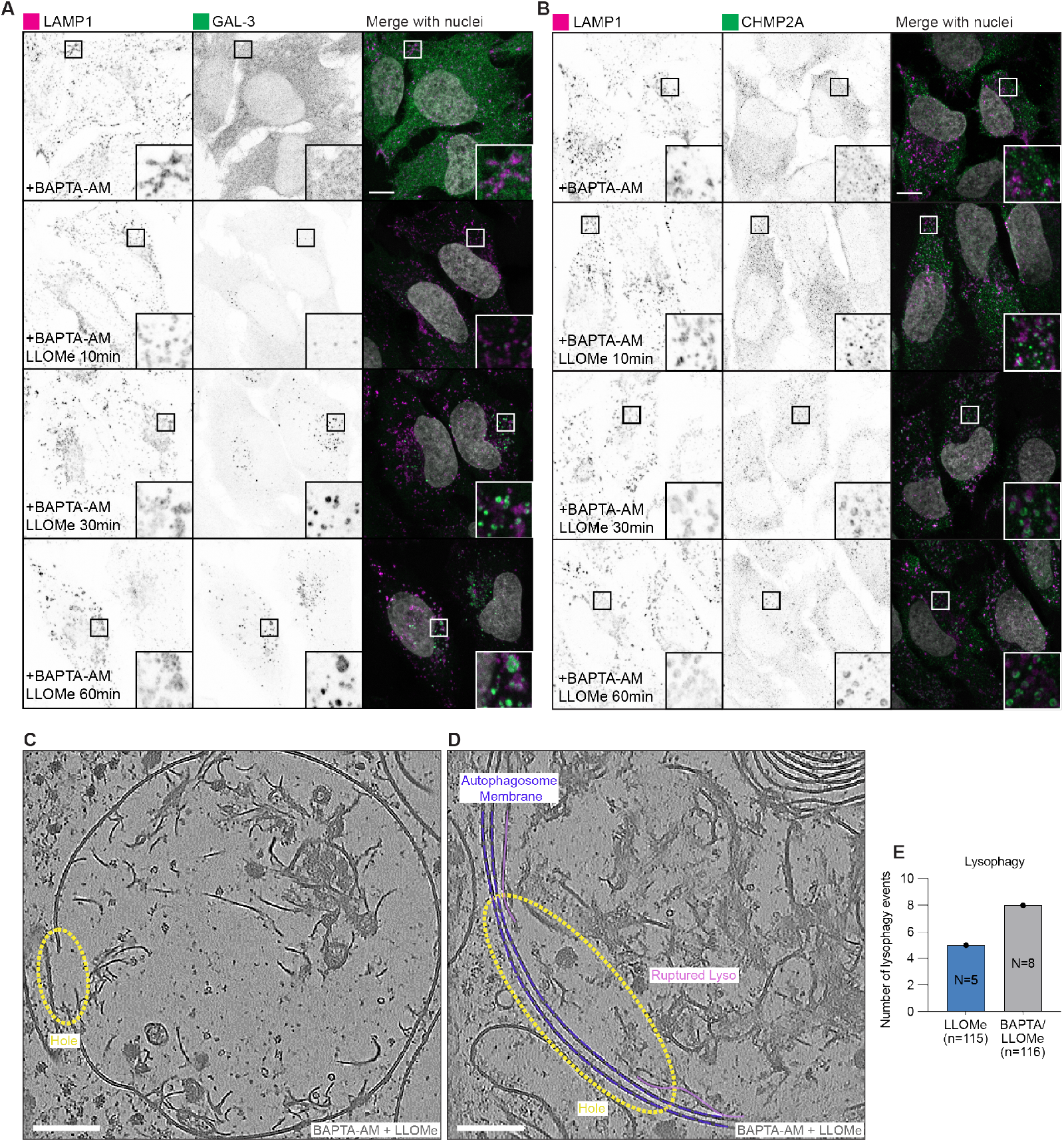
Ca2+-chelator BAPTA-AM pre-treatment leads to extensive damage. (**A**) Confocal images of immunofluorescence against LAMP1 (lysosome) and GAL-3 (lysosomal damage marker) with BAPTA-AM pre-treatment (50 µM, 30 min). Split channels and a merged image with Hoechst (nucleus) are shown. Untreated control and timepoints (10, 30, 60 min) after LLOMe treatment (0.5 mM) are indicated. (**B**) Same timepoints and treatments as in A but stained for CHMP2A. Scale for IF panels: 10 µm. (**C**) Tomographic slice of BAPTA-AM (50 µM, 30 min) LLOMe treated (0.5 mM, 60 min) lysosome with an apparent hole (circled in yellow). Scale: 100 nm. (**D**) Tomographic slice of lysosomes in BAPTA-AM+LLOMe treated cells engulfed in an autophagosome (autophagosome membrane outlined in blue, lysosomal membrane in purple). Scale: 100 nm. (**E**) Quantification of lysophagy events observed across LLOMe (N=5, n=115 tomograms) and BAPTA-AM+LLOMe datasets (N=8, n=116 tomograms).

**Fig. S4.**
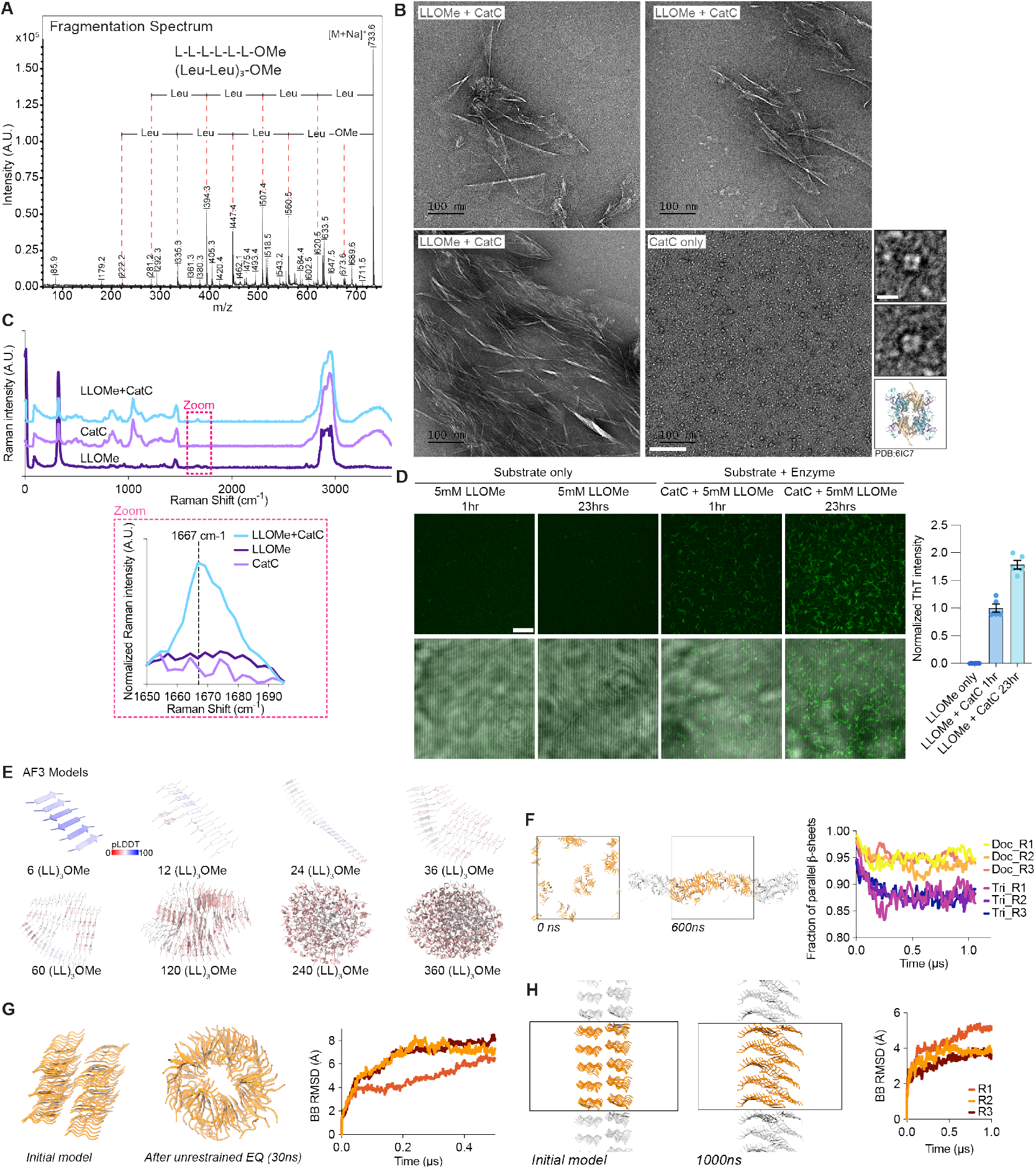
In vitro reconstitution of LLOMe formed amyloids. (**A**) Fragmentation spectrum of (LL)_3_OMe peak shown in Fig. 3A indicating the chemical composition of found species. (**B**) Negative Stain EM micrographs of LLOMe (5 mM) with CatC (5 ng/µL) and CatC alone at pH 7.5. Scale: 100 nm and 10 nm for insets. CatC atomic model (PDB:6IC7) shown for comparison. (**C**) Raman spectra of LLOMe (1 mM), CatC (1 ng/µL) and LLOMe+CatC. Background-subtracted, z-score Raman intensities are shown. (**D**) ThT staining of LLOMe amyloids without CatC present (5 mM, 1 h and 23 h) and active CatC (1 ng/µL). Quantification of ThT intensity of LLOMe only and LLOMe+CatC for 1 h and 23 h (n=5, mean ± s.e.m.). (**E**) Alphafold3 models of 6, 12, 24, 36, 60, 120, 240, 360 (LL)_3_OMe molecules, colored according to AF3 prediction confidence (pLDDT). (**F**) MD simulation of β-sheets fragments spontaneously forming fibril-like structures after 600 ns. Plot showing fraction of parallel β-sheets for pre-formed parallel trimers (Tri) or dodecamers (Doc) at 400 K in 1 µs simulation time over three repeats. (**G**) MD simulation of assembled parallel β-sheets that spontaneously form a fibril and RMSD plot over 0.5 µs to visualize ensemble stability over three repeats. (**H**) Stack of (LL)_3_OMe assemblies in sheet-like conformation initially and after 1 µs simulation time. BB RMSD plot showing fibrils remaining stable over 1 µs simulation time.

**Fig. S5.**
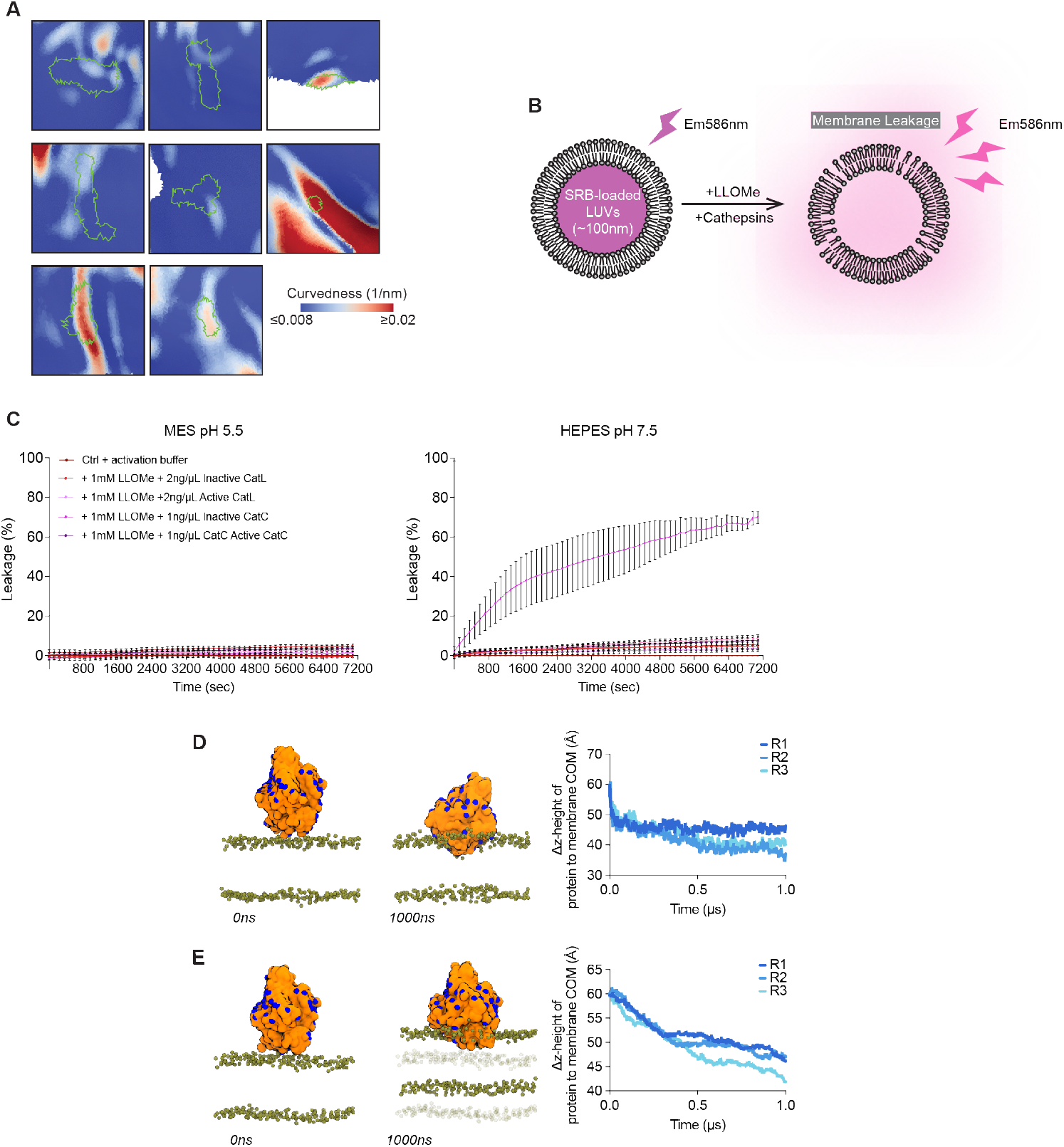
In vitro and in silico LLOMe fibrils are able to damage membranes. (**A**) Examples of the fibril associated patches (green outline) overlayed with curvedness measurements represented on lysosomal surface mesh. Scale is indicated in figure. (**B**) Schematic of liposome leakage assay mimicking lysosomal damage. SRB-fluorescence was used as readout of membrane leakage. (**C**) Percentage of leakage of liposomes at pH 5.5 (left) and pH 7.5 (right) across all conditions and time (n=3, mean ± s.e.m.). (**D**) MD simulation of LLOMe fibril (orange) freely moving in simulation box with membrane bilayer. Phosphate atoms are colored green, fibril’s nitrogen at N-termini is colored in blue. Initial fibrils position is shown vs. fibril position after 1000 ns. Plot showing z-distance between protein and membrane center of mass (COM) over three repeats. (**E**) MD simulation of LLOMe fibril (orange) fixed on distal end towards membrane. Initial position is shown vs. position after 1000 ns. Initial phosphate atom positions are shown in transparent. Plot showing z-distance between protein and membrane COM for three repeats.

**Fig. S6.**
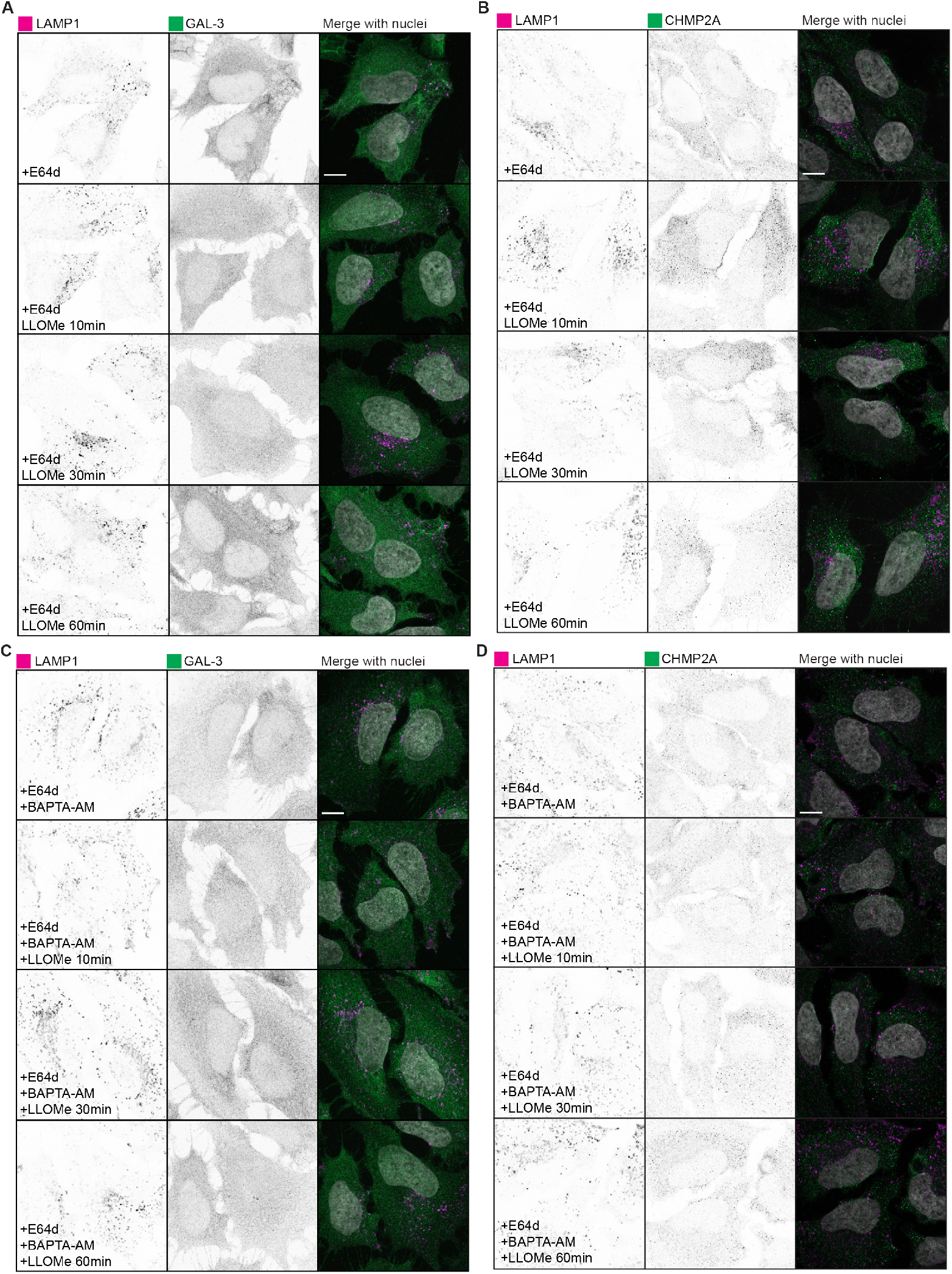
Cellular responses to E64d ±BAPTA-AM before LLOMe-treatment. (**A**) Confocal images of immunofluorescence against LAMP1 (lysosome) and GAL-3 (lysosomal damage marker) with E64d pre-treatment (100 µM, 30 min). Split channels and a merged image with Hoechst (nucleus) are shown. Untreated control and timepoints (10, 30, 60 min) after LLOMe treatment (0.5 mM) are indicated. (**B**) Same timepoints and treatments as in A but stained for CHMP2A. (**C**) Confocal images of immunofluorescence against LAMP1 (lysosome) and GAL-3 (lysosomal damage marker) with E64d (100 μM, 30 min) +BAPTA-AM (50 µM, 30 min) pre-treatment. Split channels and a merged image with Hoechst (nucleus) are shown. Timepoints (10, 30, 60 min) after LLOMe treatment (0.5 mM) are indicated. (**D**) Same timepoints and treatments as in C but stained for CHMP2A. Scale for IF panels: 10 µm.

**Fig. S7.**
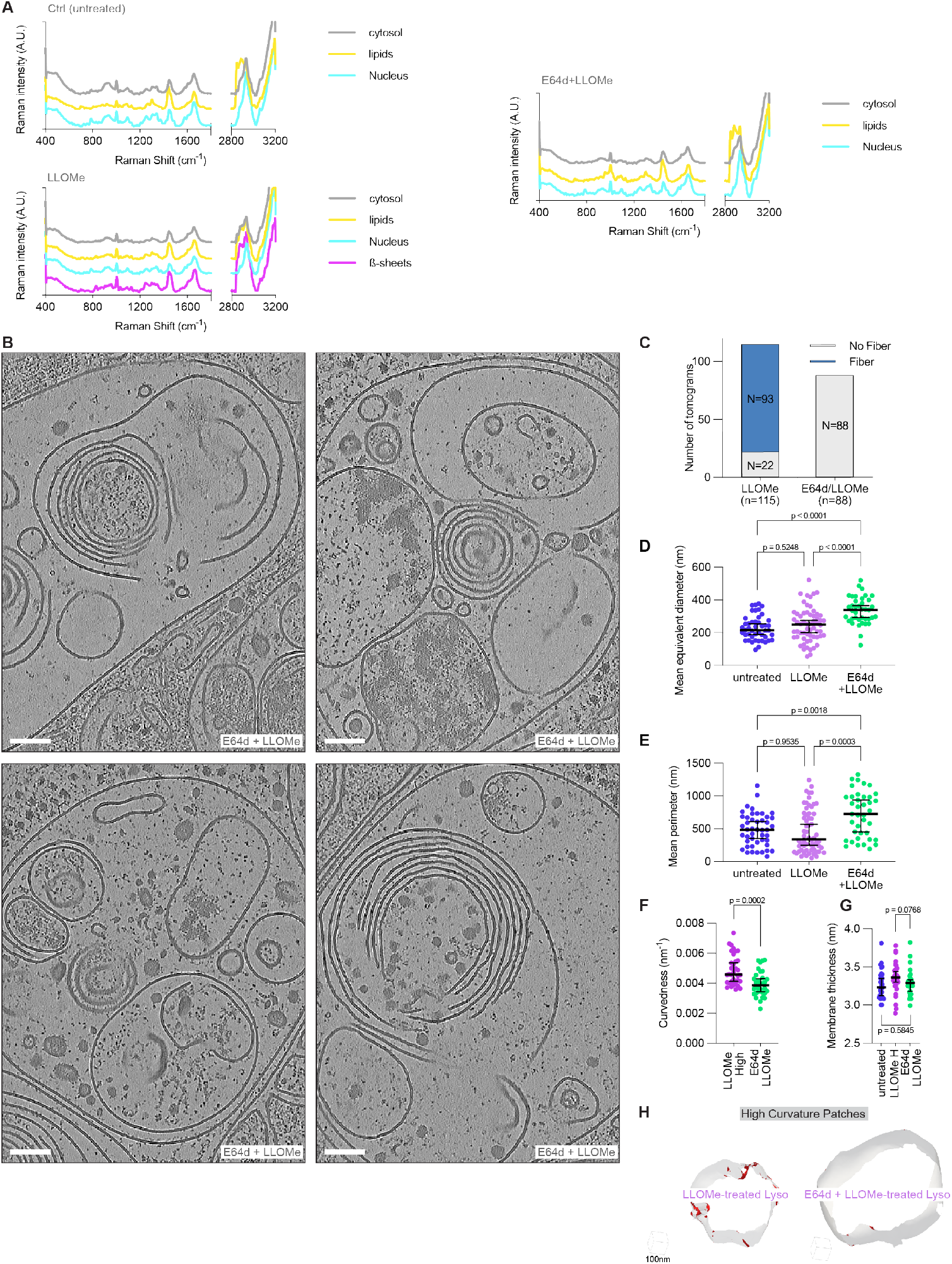
E64d negates LLOMe-amyloids by inhibiting CatC. (**A**) Raman spectra of cells shown in Fig. 4M (untreated, LLOMe, and E64+LLOMe). Spectra of nucleus, cytosol, lipids, and β-sheets-regions are visualized separately. Between 1800 cm^-1^ and 2800 cm^-1^ no peaks could be detected therefore x-axis was cut for visual purposes. (**B**) Tomographic slice of lysosomes in E64d (50 µM, 30 min) LLOMe (0.5 mM, 60 min) treated cells. Scale: 100 nm. (**C**) Quantification of tomograms with amyloid fibrils (N) inside lysosomes between with (n=88) and without (n=115) E64d pre-treatment before LLOMe. (**D**) Mean equivalent distance measurements of lysosomes from three different datasets (untreated (n=46), LLOMe (n=60), E64d+LLOMe (n=39)). (**E**) Mean perimeter measurements. Individual data points are indicated, median with 95% confidence interval drawn. Statistical test: one-way ANOVA with multiple comparisons test, p-values are indicated. (**F**) Morphometric analysis of E64d+LLOMe tomograms for general organelle curvedness. Comparison between high-damaged LLOMe lysosomes and E64d+LLOMe lysosomes. (**G**) Comparison of membrane thickness between untreated, highly-damaged LLOMe, and E64d+LLOMe lysosomes. (**H**) Lysosome membrane segmentation rendered with red highlights for areas curved above 0.021 nm^-1^.

**Movie S1**.

In situ cryo-ET tomogram of LLOMe-damaged lysosome shown in Fig. 1F (0.5 mM, 60 min).

**Movie S2**.

MD simulation rendering of fixed fibril inserting into lysosome-like membrane, shown in Fig. S5E. Simulation time: 1 µs. Fibril is colored in orange, phosphates in green. The original positions of the membrane phosphates are shown as transparent spheres.

**Movie S3**.

MD simulation rendering of fibril inserted in membrane during equilibration phase, creating holes in both sides of the lysosome-like membrane, shown in Fig. 4I. Simulation time: 1 µs. Fibril is colored in orange, phosphates in green, and water molecules are colored in licorice. Na^+^ (yellow) and Cl^-^ (lightblue) ions are displayed as spheres.

